# Molecular properties of RmlT, a wall teichoic acid rhamnosyltransferase that modulates virulence in *Listeria monocytogenes*

**DOI:** 10.1101/2024.01.23.574818

**Authors:** Ricardo Monteiro, Tatiana B. Cereija, Rita Pombinho, Thijs Voskuilen, Jeroen D.C. Codée, Sandra Sousa, João H. Morais-Cabral, Didier Cabanes

**Affiliations:** i3S – Instituto de Investigação e Inovação em Saúde, Universidade do Porto, Porto, Portugal; IBMC – Instituto de Biologia Molecular e Celular, Universidade do Porto, Porto, Portugal; Leiden Institute of Chemistry, Leiden University, The Netherlands

**Author notes:** These authors contributed equally (Ricardo Monteiro, Tatiana B. Cereija). These authors share last authorship (João H. Morais-Cabral, Didier Cabanes).

## Abstract

Wall teichoic acids (WTAs) from the major Gram-positive foodborne pathogen *Listeria monocytogenes* are peptidoglycan-associated glycopolymers decorated by monosaccharides that, while not essential for bacterial growth, are required for bacterial virulence and resistance to antimicrobials. Here we report the structure and function of a bacterial WTAs rhamnosyltransferase, RmlT, strictly required for *L. monocytogenes* WTAs rhamnosylation. In particular, we demonstrated that RmlT transfers rhamnose from dTDP-L-rhamnose to naked WTAs, and that specificity towards TDP-rhamnose is not determined by its binding affinity. Structures of RmlT with and without its substrates showed that this enzyme is a dimer, revealed the residues responsible for interaction with the substrates and that the catalytic residue pre-orients the acceptor substrate towards the nucleophilic attack to the sugar. Additionally, the structures provided indications for two potential interaction pathways for the long WTAs on the surface of RmlT. Finally, we confirmed that WTAs glycosyltransferases are promising targets for next-generation strategies against Gram-positive pathogens by showing that inactivation of the RmlT catalytic activity results in a decreased infection *in vivo*.

## Introduction

Infections caused by Gram-positive bacteria remain a major public health burden and a main challenge in medicine. We witness the emergence of not only new pathogens but also new bacterial strains with altered virulence properties and/or increased antimicrobial resistance [1]. Among them, *Listeria monocytogenes* is the Gram-positive foodborne pathogen that accounts for the highest proportion of hospitalisation cases and number of deaths due to food poisoning in Europe, with emergence of antibiotic resistances [1, 2]. The cell wall of *L. monocytogenes* and other Gram-positive bacteria is a thick mesh-like complex that maintains cellular integrity. It is composed of three major constituents, namely the peptidoglycan backbone, anionic teichoic acid polymers and wall-associated proteins. Teichoic acids are the vast majority of cell wall carbohydrates, either associated to the peptidoglycan (wall teichoic acids or WTAs) or linked via a lipid anchor to the membrane in gram-positive bacteria (lipoteichoic acids or LTAs) [3]. While the basic structure of LTAs remains relatively conserved among different Gram-positive species and serovars [4–8], WTAs display significant structural and constitutional diversity, as well as a variety of glycosylation patterns, resulting in different fitness outcomes such as biofilm formation, virulence capacity and resistance to bacteriophages and antimicrobials [4, 9–21]. WTAs are complex glycopolymers made of 20 to 30 repeating units of ribitol-phosphate (RboP). In *Staphylococcus aureus*, the polymeric RboP backbone of WTAs is modified through the addition of *N*-acetylglucosamine (GlcNAc) moieties by different glycosyltransferases, including TarM, TarS, and TarP, which have different specificities. While TarM is responsible for *α*-GlcNAc glycosylation at position *C*-4 of the ribitol unit, TarS and TarP are required for the *β*-GlcNAc glycosylation at positions *C*-4 and *C*-3, respectively [22–24]. Interestingly, all these individual GlcNAc additions to the poly-RboP chain have a dramatic impact on pathogen-host interactions [25]. The molecular properties of the *S. aureus* WTAs glycosyltransferases have been characterized [24–27].

*L. monocytogenes* WTAs are grouped in two subtypes: i) Type I WTAs, made of repeating units of 1,5-phosphodiester-linked ribitol-phosphate glycosylated with GlcNAc at position *C*-2 and rhamnose at position *C*-4 in serovar 1/2, with GlcNAc at position *C*-4 in serovar 3, or non-glycosylated in serovar 7; and ii) Type II WTAs integrate a GlcNAc moiety into the repeating unit at positions *C*-2 in serovars 4a, 4c, 5, 6b or at *C*-4 in serovars 4b, 4d, 4e, 4h and 6a in addition to other modifications with glucose, galactose or GlcNAc moieties, or with acetyl groups [4, 20].

We previously performed the first genome-wide transcriptome of a bacterial pathogen infecting a host organism using *L. monocytogenes* EGDe (serovar 1/2a) strain as model pathogen [28]. This study revealed that genes responsible for the biosynthesis of dTDP-L-rhamnose (*rmlABCD*) and the transfer of the rhamnose moiety to WTAs (*rmlT*) are highly expressed during *in vivo* infection [28]. Importantly, we showed that WTAs rhamnosylation is required for the proper association of major virulence factors to the bacterial surface [14], and for resistance to antimicrobial peptides [13] and antibiotics [16]. In addition, we showed that WTAs decoration with rhamnose is strictly dependent on the rhamnosyltransferase RmlT [13]. Altogether, our previous data revealed that, although dispensable for bacterial growth, RmlT has a significant impact on bacterial virulence and antimicrobial resistance.

Here, we explored the molecular properties of RmlT. Using a combination of structural and biochemical approaches, we solved the structure of RmlT, uncovered its oligomeric organization, identified the molecular determinants of donor-substrate binding and specificity, and show that inactivation of RmlT catalytic activity can significantly reduce bacterial virulence, confirming this rhamnosyltransferase as a promising target for new anti-virulence strategies.

## Results

### Enzymatic activity of the *L. monocytogenes* rhamnosyltransferase RmlT

We previously showed that RmlT is required for WTAs rhamnosylation in *L. monocytogenes* [13]. To confirm that RmlT is indeed capable of transferring rhamnose to WTAs, we set-up an *in vitro* enzymatic assay using dTDP-L-rhamnose, now on referred as TDP-rhamnose, as donor substrate and naked WTAs as acceptor substrate. Naked WTAs were isolated from *L. monocytogenes* EGDe Δ*rmlT*Δ*lmo1079* strain that lacks the ability to produce both RmlT and Lmo1079, respectively reported to be involved in the addition of rhamnose and GlcNAc moieties to WTAs [13, 29]. RmlT enzymatic activity was evaluated by analysing in parallel the reaction products: decorated WTAs and TDP.

Decoration of WTAs was analyzed by electrophoretic mobility shift assay. After incubation with TDP-rhamnose and purified RmlT, WTAs showed a change in migration, with bands running higher in a native gel and displaying a more diffuse appearance compatible with glycosylated material (Figure 1A). The generation of TDP by RmlT when transferring rhamnose to WTAs was quantified indirectly using a coupled functional assay that measures the amount of free phosphate generated by a phosphatase from the TDP produced by RmlT activity (Figure 1B). The amount of phosphate detected in the reaction mixture containing TDP-rhamnose, WTAs and RmlT was significantly higher than in the reaction mixture lacking RmlT. The generation of TDP only happens when naked WTAs is present, showing that RmlT is not capable of hydrolysing TDP-rhamnose to TDP and rhamnose in the absence of the acceptor substrate WTAs. Together, these experiments strongly indicate that RmlT transfers rhamnose from TDP-rhamnose to naked WTAs, producing TDP and rhamnosylated WTAs.

**Figure 1:**
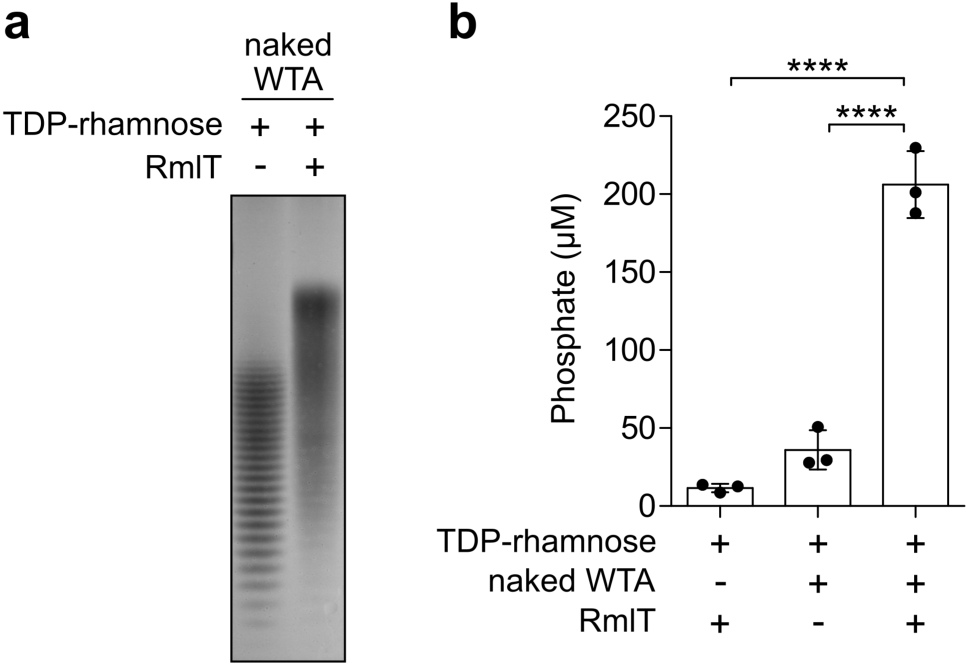
RmlT transfers rhamnose from TDP-rhamnose to naked WTAs. a) Electrophoretic mobility shift assay of the reaction mixture containing naked WTAs and TDP-rhamnose, in the absence (−) and presence (+) of RmlT. The individual bands observed in each lane correspond to different WTAs chains. **b)** Coupled functional assay quantifies the amount of phosphate released after hydrolysis of TDP molecules generated by RmlT. In both assays reactions ran for 60 minutes with 5 mM WTAs, 1 mM TDP-rhamnose and 10 µM RmlT. Mean ± SD (n = 3) and individual measurements are shown; one-way ANOVA; **** *p* < 0.0001.

### Structure of RmlT

To gain insights into the molecular properties of *L. monocytogenes* RmlT, we determined the crystal structures of the full-length protein in different states at resolutions varying between 2.2 and 2.5 Å (Table 1). All the structures described below include multiple copies in the asymmetric unit. As assessed from the electron-density map, different copies display varying quality in different parts of the protein. Therefore, figures below show the best locally-defined protein copy, with its chain ID indicated in the legend. We also indicate the chain ID of the copy(ies) that is(are) globally the best defined (lowest average RSRZ value) in Table 1 and Supplementary Table 1.

**Table 1.** Data collection, phasing and refinement statistics.

|  | Apo<br>SeMet-RmlT | Apo RmlT/<br>HEPES | RmlT/<br>UDP-glucose | RmlT/TDP-rhamnose | RmlT<sub>D198A</sub>/TDP<br>-rhamnose | RmlT/<br>3-RboP | RmlT/<br>4-RboP |
| --- | --- | --- | --- | --- | --- | --- | --- |
| PDB ID | 8BZ4 | 8BZ5 | 8BZ6 | 8BZ7 | 8BZ8 | 9GZJ | 9GZK |
| Diffraction data |  |  |  |  |  |  |  |
| Space group | P 2<sub>1</sub> 2<sub>1</sub> 2<sub>1</sub> | P 1 2<sub>1</sub> 1 | P 1 2<sub>1</sub> 1 | P 1 2<sub>1</sub> 1 | P 1 2<sub>1</sub> 1 | P 2<sub>1</sub> 2<sub>1</sub> 2<sub>1</sub> | P 1 2<sub>1</sub> 1 |
| Cell dimensions |  |  |  |  |  |  |  |
| *a*, *b*, *c* (Å) | 83.4, 103.7,<br>178.4 | 82.1, 178.4,<br>92.2 | 85.5, 291.1,<br>89.8 | 87.0, 291.3,<br>92.4 | 87.4, 292.4,<br>93.1 | 83.2, 103.4,<br>178.6 | 86.3, 290.8, 91.0 |
| *α*, *β*, *γ* (°) | 90.0, 90.0, 90.0 | 90.0, 93.5, 90.0 | 90.0, 100.1,<br>90.0 | 90.0, 100.4,<br>90.0 | 90.0, 100.5,<br>90.0 | 90.0, 90.0,<br>90.0 | 90.0, 100.2, 90.0 |
| Resolution (Å) | 49.8 – 2.52 | 48.1 – 2.30 | 49.3 – 2.40 | 50.0 – 2.20 | 49.3 – 2.52 | 48.4 – 2.34 | 49.0 – 2.52 |
| *R*<sub>merge</sub><sup>*</sup> | 0.210 (1.735) | 0.112 (1.295) | 0.103 (1.560) | 0.059 (1.087) | 0.113 (1.394) | 0.079 (1.363) | 0.071 (1.053) |
| *I* / *σI*<sup>*</sup> | 9.8 (1.5) | 10.7 (1.4) | 12.2 (1.3) | 14.0 (1.2) | 12.9 (1.6) | 16.2 (1.2) | 13.2 (1.4) |
| Completeness<sup>*</sup> (%) | 100 (100) | 99.2 (98.4) | 99.1 (97.7) | 99.8 (99.7) | 100 (100) | 99.9 (98.7) | 99.5 (99.8) |
| Redundancy<sup>*</sup> | 13.6 (13.3) | 7.0 (7.2) | 7.2 (7.3) | 3.8 (3.6) | 7.1 (7.1) | 6.6 (6.0) | 3.5 (3.5) |
| Refinement |  |  |  |  |  |  |  |
| Resolution (Å) | 49.8 – 2.52 | 48.1 – 2.30 | 44.2 – 2.40 | 50.0 – 2.20 | 45.8 – 2.52 | 41.6 – 2.34 | 42.9 – 2.52 |
| No. reflections | 53,011 | 115,974 | 166,253 | 227,111 | 153,926 | 65,950 | 147,192 |
| *R*<sub>work</sub> / *R*<sub>free</sub> | 0.219 / 0.251 | 0.193 / 0.220 | 0.195 / 0.235 | 0.187 / 0.215 | 0.191 / 0.228 | 0.192 / 0.226 | 0.198 / 0.232 |
| Total No. atoms | 9,848 | 20,336 | 30,490 | 30,744 | 30,233 | 9,962 | 29,353 |
| Total No. water<br>molecules | 241 | 502 | 674 | 1,193 | 673 | 282 | 334 |
| R.m.s deviations |  |  |  |  |  |  |  |
| Bond lengths (Å) | 0.003 | 0.004 | 0.004 | 0.005 | 0.002 | 0.003 | 0.002 |
| Bond angles (°) | 0.521 | 0.586 | 0.581 | 0.670 | 0.470 | 0.552 | 0.424 |
| Ramachandran plot |  |  |  |  |  |  |  |
| Favored (%) | 95 | 96 | 96 | 97 | 97 | 96.58 | 95.69 |
| Allowed (%) | 5 | 4 | 4 | 3 | 3 | 3.42 | 4.31 |
| Outliers (%) | 0 | 0 | 0 | 0 | 0 | 0 | 0 |
| Ligand | - | - | UDP-glucose | TDP-rhamnose | TDP-rhamnose | 3-RboP | 4-RboP |
| Molecules in the<br>asymmetric unit | 2 | 4 | 6 | 6 | 6 | 2 | 6 |
| Best defined chain<br>(Mean RSRZ) | A (0.04) | B (0.03) | E (0.15) | A (0.12) | A, C, D (0.04) | A (-0.32) | A, C, D, E (-0.17) |
| Residues<br>NOT<br>modeled<br>(per<br>chain) | A: M1-K17; B:<br>M1-K20, E31-E33, G60-T65,<br>K82-G90, G227-F234. | A: M1-N15, A228-S232; B: M1- K16,<br>A228-F234; C: M1-N15; D: M1-K16;<br>S87-W89; G227-S232. | All chains:<br>M1-K17; B:<br>also missing<br>L623. | A: M1-K17, A228-S232; B: M1-K17,<br>N230-F234; C: M1-K17, N230-G235; D:<br>M1- K16, A228-G235;<br>E and F: M1-K17,<br>G227-F234. | A, C, D: M1-K17,<br>N230-S232; B:<br>M1-K17, A228-S232; E, F: M1-K17, G227-S233. | A: G0-N15;<br>B: G0-D18,<br>T30-G34,<br>D58-Y66,<br>K82-G90,<br>G227-F234,<br>L623. | A: G0-K16, A228-F234; B: G0-K17,<br>A228-S232; C and<br>D: G0-K17, G227-S233; E: G0-K17,<br>G227-S231; F: G0-K17, T62-T75,<br>A228-F234, L536-T545, L565-577,<br>L596-V607, I619-L623. |

The ∼2.5Å crystal structure of RmlT in the apo state (PDB code: 8BZ4) presents the characteristic architecture of other WTAs glycosyltransferases like the *S. aureus* TarS and TarP [24, 27], with a catalytic region at the N-terminus and an oligomerization region at the C-terminus (Figure 2A and Supplementary Figures 1A and 2). Running from the N- to the C-terminus, RmlT shows a catalytic subdomain strongly connected to a helical subdomain, a large oligomerization subdomain and a turret subdomain (Figure 2A). Comparison between RmlT, TarS and TarP (all classified as members of the GT2 family of glycosyltransferases in CAZy, (http://www.cazy.org/), showed strong similarities at the level of the catalytic subdomains of the three enzymes (Figure 2B), while the helical, oligomerization and turret subdomains of RmlT are very similar to the equivalent regions in TarS (Supplementary Figure 1A) [27]. Interestingly, it has been shown that the oligomerization and turret domains of TarS stand apart from the catalytic domain, adopting different dispositions that are suggestive of flexibility [27]. In contrast, the oligomerization domain of RmlT lies over the catalytic domain, establishing an extensive interaction surface (∼480 Å^2^) (Figure 2A and Supplementary Figure 1A). In addition, the connection between the helical and oligomerization subdomains is established in TarS by a 4 amino acid linker (residues 349-352) [27] while in RmlT three linkers (residues 340-342, 368-388 and 406-410) run between the subdomains (Supplementary Figure 1B).

**Figure 2:**
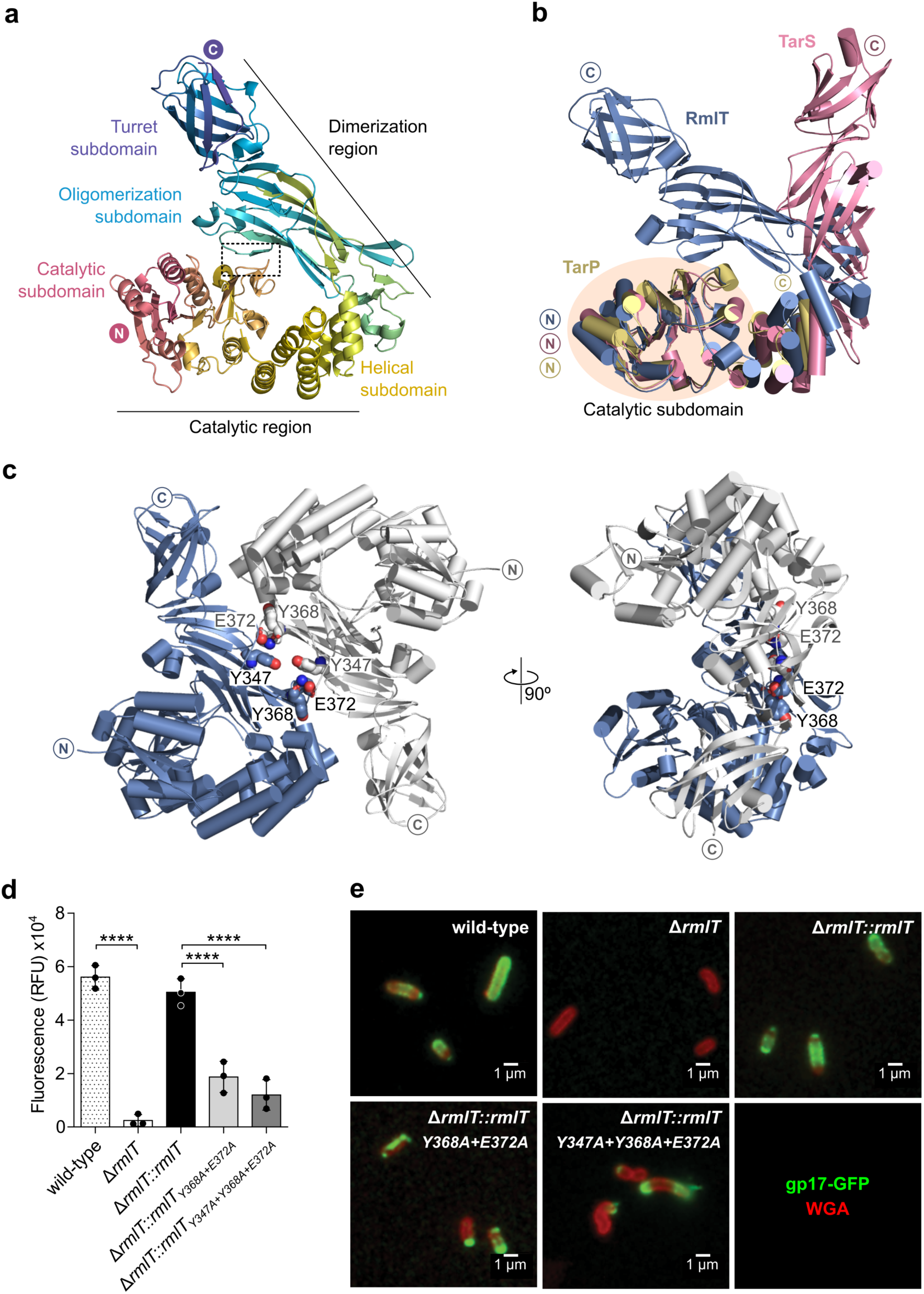
Structure of *L. monocytogenes* RmlT. **a)** Cartoon representation of a subunit of RmlT (PDB code: 8BZ4; chain A) coloured from pink to purple, with subdomains and N and C termini indicated. Interaction between catalytic and oligomerization subdomains is indicated by dashed rectangle. **b)** Superposition of RmlT (blue, PDB code: 8BZ4, chain A) and *S. aureus* TarS [27] (pink, PDB code: 5TZ8) and TarP [24] (yellow, PDB code: 6H1J) through catalytic domain (wheat ellipse) Cα atoms in helices h5, h6 and h11 (RmlT: residues 91-101, 118-130, 196-206; TarS: residues 72-82, 99-111,176-186; TarP: residues 73-83; 100-112; 179-189) with RMSD values of 1.185 Å (TarS) and 0.886 Å (TarP). **c)** Two views of RmlT dimer showing subunits coloured blue and white and dimerization interface residues Y347, Y368 and E372 as spheres. **d)** Quantification of fluorescence associated with gp17-GFP bound to rhamnose at the surface of the *L. monocytogenes* wild-type EGDe strain and mutant strains EGDe Δ*rmlT::rmlT,* EGDe Δ*rmlT::rmlT_Y368A+E372A_* and EGDe Δ*rmlT::rmlT_Y347A+Y368A+E372A_*. Mean ± SD (n = 3) and individual measurements are shown; one-way ANOVA, **** = *p* < 0.0001. **e)** Fluorescence microscopy images of *L. monocytogenes* cells stained with wheat germ agglutinin (WGA, red) and gp17-GFP (green) which binds to rhamnose at the bacterial surface.

Contrasting with Tar proteins that form trimers (Supplementary Figure 1C) [27], RmlT eluted as a dimer in a size-exclusion chromatography column (Supplementary Figure 3). Moreover, RmlT assembles as a dimer in the crystal with the oligomerization subdomains interacting in an antiparallel arrangement and positioning the catalytic subdomains at opposite corners of a rectangular shape (Figure 2C). To assess the dimeric architecture of RmlT, we generated structure-guided mutants (RmlT_Y368A+E372A_ and RmlT_Y347A+Y368A+E372A_) that aimed at disrupting the dimerization interface (Figure 2C). Attempts to express and purify mutant proteins resulted in aggregated protein. Nevertheless, after optimization of expression [30], we successfully purified a small amount of both mutant proteins. By size exclusion chromatography, we showed that both mutant proteins elute as monomers, confirming that mutated residues are part of the dimer interface and that wild type RmlT is a dimer in solution (Supplementary Figure 3). Interestingly, mutations at the dimerization interface did not affect the catalytic activity of RmlT *in vitro* (Supplementary Figure 3C). The impact of the mutations at the RmlT dimerization interface was also tested *in vivo* using a bacteriophage rhamnose binding protein (gp17), that specifically binds to rhamnose at the *L. monocytogenes* bacterial surface, fused to GFP (gp17-GFP) [24]. Exponentially growing bacteria were incubated with gp17-GFP and fluorescence associated with the bacterial cell wall staining was quantified in a microtiter plate format and visualized by fluorescence microscopy (Figure 2D-E). Deletion of *rmlT* in the *L. monocytogenes* EGDe strain resulted in a very significant decrease in the fluorescence intensity to background levels, which was restored by complementation with *rmlT*, confirming the role of RmlT in the rhamnosylation of the *L. monocytogenes* cell surface. Complementation with the *rmlT* dimerization interface mutants significantly reduced, but did not abolish, fluorescence intensity (Figure 2D). Fluorescence microscopy showed that the surface of bacteria expressing the dimerization mutants (EGDe Δ*rmlT*::*rmlT*_Y368A+E372A_ and EGDe *ΔrmlT*::*rmlT*_Y347A+Y368A+E372A_) is still partially rhamnosylated, consistent with their activity *in vitro*, but exhibited an altered pattern, with staining restricted to the poles (Figure 2E). The mechanism underlying this change in staining pattern is not known, but altogether these data demonstrate that RmlT is a dimer in solution and that changes in dimer formation impact its physiological function.

### RmlT active site

The structure of RmlT with bound TDP-rhamnose (PDB code: 8BZ7) was solved at 2.2 Å with 6 subunit copies in the asymmetric unit (Figure 3A, Table 1 and Supplementary Table 1). The electron-density for the ligand is clear and it was straightforward to build the nucleoside moiety in the catalytic subdomain (Supplementary Figures 4A-B and 5). However, there is a lack of detail in the electron-density for the sugar. The apparent spatial disorder of the sugar is evident in a superposition of the 6 copies of RmlT/TDP-rhamnose, while the nucleotide base and deoxyribose groups are unchanged between copies (Supplementary Figure 6A).

**Figure 3:**
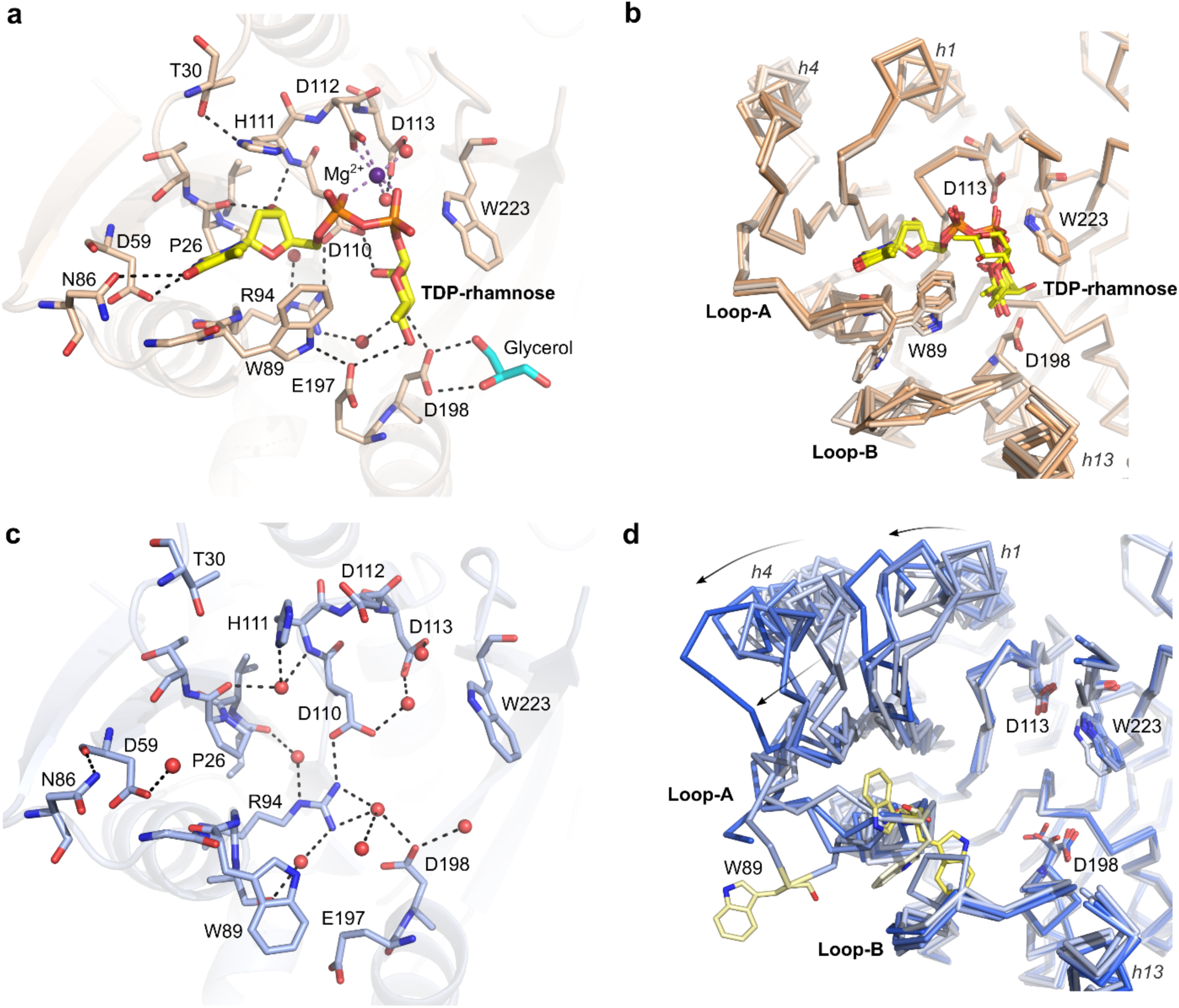
Active site of RmlT. **a)** Cartoon representation of the active-site region of RmlT with TDP-rhamnose and interacting residues represented as sticks (PDB code: 8BZ7, chain D). Glycerol molecule used as cryo-protectant was found interacting with D198, likely mimicking a ribitol molecule. Water molecules and Mg^2+^ are shown as red and purple spheres, respectively. Hydrogen bonds and metal coordination are shown as black or light purple dashed lines. **b)** Superposed active-sites (through residues 110 to 114) of RmlT/TDP-rhamnose copies (chains A to F) present in the asymmetric unit, coloured in different shades of wheat (PDB code: 8BZ7). TDP-rhamnose and some substrate-interacting residues are shown as sticks. RMSD relative to chain D: 0.034 Å (chain A), 0.046 Å (chain B), 0.027 Å (chain C), 0.042 Å (chain E) and 0.060 Å (chain F). **c)** Cartoon representation of the active-site region of the apo RmlT (chain A) with donor substrate interaction residues represented as sticks (PDB code: 8BZ4). **d)** Superposed active-sites (through residues 110 to 114) of apo RmlT copies present in the asymmetric unit of two different crystal forms, coloured in different shades of blue (PDB codes: 8BZ4 and 8BZ5). RMSD relative to chain A of 8BZ4: 0.966 Å (8BZ4 chain B), 0.319 Å (8BZ5 chain A), 0.229 Å (8BZ5 chain B), 0.070 Å (8BZ5 chain C) and 0.807 Å (8BZ5 chain D). Some substrate-interacting residues are represented as sticks with W89 in yellow. Loops A and B and helices *h*1, *h*4 and *h*13 are indicated. Positional variability of *h*1 and *h*4 is represented by arrows.

Comparison of catalytic subdomains of RmlT/TDP-rhamnose, TarP/UDP-GlcNAc (PDB code: 6H2N) [24] and TarS/UDP-GlcNAc (PDB code: 5TZE) [27] showed that many of the molecular interactions established by the nucleoside moiety of the donor substrate involve conserved residues (Supplementary Figures 2 and 6). In particular, in RmlT (Figure 3A) the nucleotide base is hydrogen bonded to D59 and N86, while the deoxyribose hydroxyl interacts with the main-chain carbonyl of the conserved P26 and amino group of H111. In addition, D112 and D113 together with the phosphate groups of the nucleotide are involved in the coordination of a single Mg^2+^. D112 coordinates the cation directly while D113 interacts *via* two water molecules, just like in the other two enzymes (Supplementary Figure 6). The conserved residue D110 lines the bottom of the binding pocket by interacting with the also conserved R94. Of the interactions between the rhamnose and the protein, those involving E197 and D198 standout as these residues are conserved in TarP and TarS and are considered to be crucial for catalysis (Figure 3A and supplementary Figure 6) [24, 27]. In addition, two non-conserved tryptophans (W89 and W223) also interact with the sugar. The position of W89 varies between RmlT copies (Figure 3B) but that of W223 is invariant and is within hydrogen bond distance of the sugar oxygen (*O*1) (Figure 3B), raising the hypothesis that it contributes to the polarization of the anomeric carbon promoting the catalytic reaction.

In addition to the structures of apo-RmlT described above, with 2 copies in the asymmetric unit, and of RmlT/TDP-rhamnose complex (6 copies), we also determined another RmlT apo-structure with 4 copies per asymmetric unit (PDB code: 8BZ5) at a slightly higher resolution (2.3 Å) (Figure 3C, Table 1 and Supplementary Table 1). Globally, there are only minor changes between bound and unbound structures. Comparison between the donor binding sites of RmlT with and without TDP-rhamnose showed a widening of the binding site in the absence of the nucleotide (Supplementary Figure 7). The availability of multiple protein copies due to non-crystallographic symmetry and different crystal forms allowed us to explore in more detail the conformational space sampled in the two states. In the nucleotide-bound subunits there is little structural variability in the catalytic subdomain (Figure 3B). In contrast, we observed large spatial variability in the apo-structures (Figures 3D), likely to reflect local structural freedom due to the absence of stabilizing interactions with the ligand. In particular, the major structural variability in the catalytic subdomain occurs in two loops, loops A (residues 86-90) and B (residues 190-195). In one of the apo-protein copies, loop A (residues 86-90) flips away from the binding site and W89 (already described above as interacting with the rhamnose in the RmlT complex) points towards the bulk solvent.

To evaluate the functional impact of some of the residues in the catalytic subdomain, we generated single-point alanine mutants at the active-site residues E197, D198, W89 and W223 (Figure 3A). Recombinant RmlT mutants were incubated with naked WTAs and TDP-rhamnose, and the reaction was quantified by the free-phosphate coupled functional assay (Figure 4A). As expected, mutation of E197 or D198 resulted in a complete loss of *in vitro* activity, fitting with their proposed role in catalysis. W223A had wild-type levels of activity, showing that this residue is not important for the catalytic reaction. In contrast, W89A resulted in a loss of function, supporting the idea that W89 has a role in the proper positioning of the sugar.

**Figure 4:**
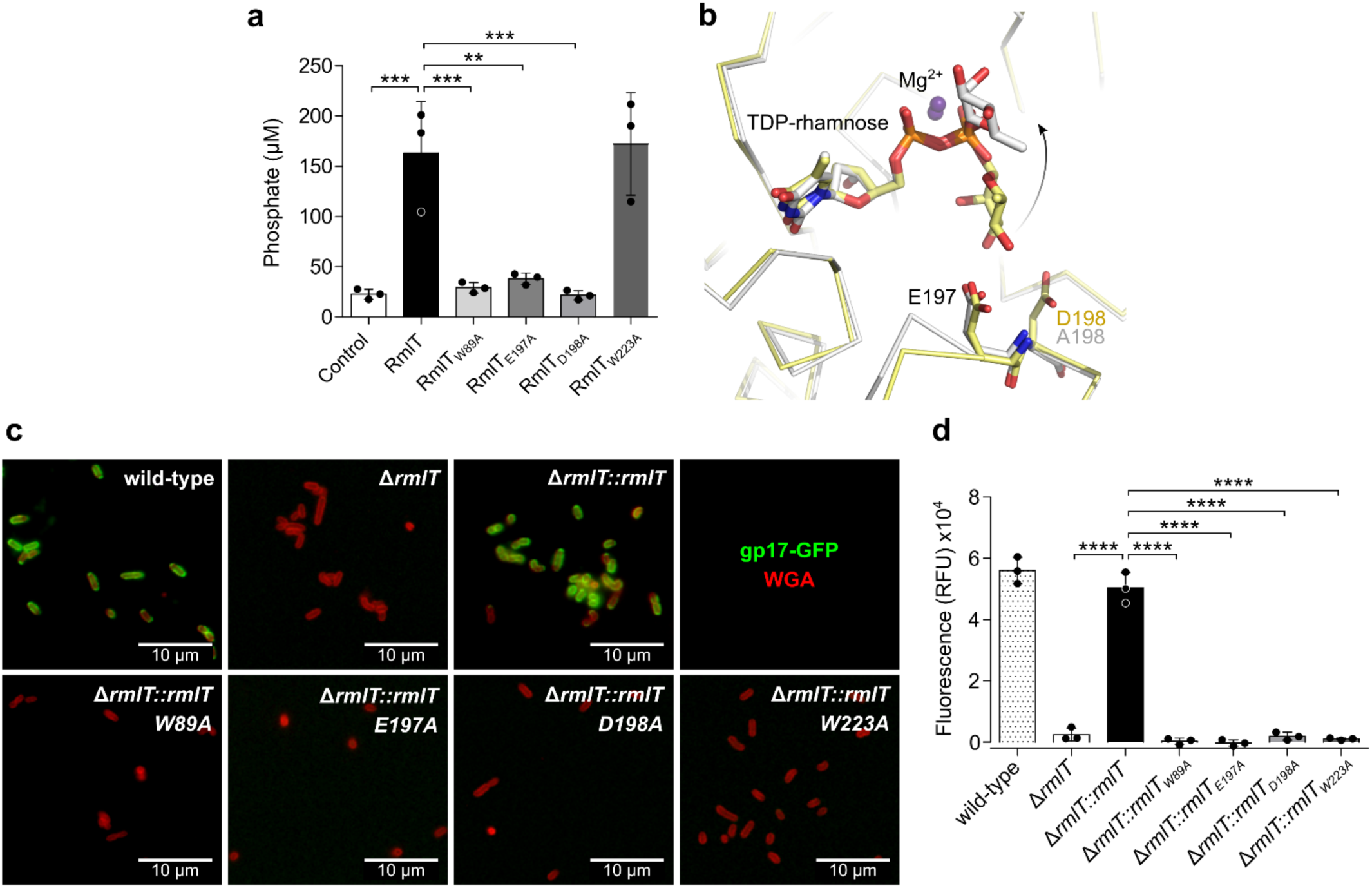
Impact of active-site residues in RmlT activity. **a)** Quantification of the amount of phosphate released from the product generated by RmlT, RmlT_W89A_, RmlT_E197A_, RmlT_D198A_ and RmlT_W223A_. Phosphate was quantified after 30 minutes reaction from mixtures containing 1 mM naked WTAs, 200 μM TDP-rhamnose and 2 μM protein. **b)** Superposition of active-sites of RmlT (light yellow, PDB code: 8BZ7; chain A) and RmlT_D198A_ (grey, PDB code: 8BZ8; chain F) in complex with TDP-rhamnose through Cα of residues 110, 112-113 and 197-198 (Cα RMSD: 0.245 Å). Apparent change in the position of rhamnose moiety is indicated by arrow. **c)** Impact of mutations in the active site of RmlT on *L. monocytogenes* surface rhamnosylation. Fluorescence microscopy analysis of *L. monocytogenes* strains EGDe wild-type, EGDe Δ*rmlT* and EGDe Δ*rmlT* complemented with *rmlT*, *rmlT_W89A_*, *rmlT_E197A_*, *rmlT_D198A_* or *rmlT_W223A_*. Bacterial cells and the rhamnose at the bacterial surface were stained with wheat germ agglutinin (WGA, red) and gp17-GFP rhamnose binding protein (green), respectively. **d)** Quantification of the fluorescence associated to gp17-GFP bound to rhamnose at the surface of the strains in c). Mean ± SD (n = 3) and individual measurements are shown; one-way ANOVA; ** *p* < 0.01, *** *p* < 0.001, **** *p* < 0.0001.

To explore further the role of D198, which in TarS and TarP has been proposed to promote the nucleophilic attack of the ribitol [24, 27], we determined the ∼2.5 Å structure of the mutant RmlT_D198A_ in complex with TDP-rhamnose (PDB code: 8BZ8) (Figure 4B, Table 1 and supplementary Table 1). The 6 copies of the protein in the asymmetric unit are occupied by nucleotide and are for the most part identical to the structures of wild-type RmlT (Supplementary Figures 4C-D, 5B and 8). In the RmlT_D198A_ mutant structure the sugar is even less defined in the electron-density and appear to have moved away in all 6 copies of the asymmetric unit from its position in the wild type protein (Figure 4B and Supplementary Figures 4C, 4D, 6B and 9). This suggests that D198 has a role in positioning the sugar, in agreement with the observation of contacts with the rhamnose in the RmlT/TDP-rhamnose structure.

We also evaluated *in vivo* the impact of mutants for residues in the catalytic subdomain. The *L. monocytogenes* EGDe Δ*rmlT* strain was complemented with wild-type RmlT or RmlT mutants (RmlT_W89A,_ RmlT_E197A_, RmlT_D198A_, and RmlT_W223A_), and the effect of each mutation was analyzed using the gp17-GFP rhamnose binding protein. Mutations of W89, E197 and D198 completely abolished rhamnose from the bacterial surface (Figures 4C-D), confirming their importance for RmlT activity. Surprisingly, W223A also induced the absence of surface rhamnosylation, which suggests that the mutation affected a property of RmlT that is specific to its physiological function *in vivo* other than catalytic reaction.

### RmlT specificity for TDP-rhamnose is not determined by ligand affinity

The lack of strong contacts between rhamnose and RmlT residues raised the possibility that the enzyme is not selective relative to the donor substrate sugar. To explore this hypothesis, we measured RmlT activity using the free-phosphate coupled functional assay in the presence of naked WTAs and the nucleotides TDP-rhamnose, TDP-glucose, UDP-glucose and UDP-GlcNAc as well as L-rhamnose. The amount of phosphate detected was higher with TDP-rhamnose relative to all other donor substrates tested, demonstrating a catalytic preference of RmlT for this nucleotide (Figure 5A).

**Figure 5:**
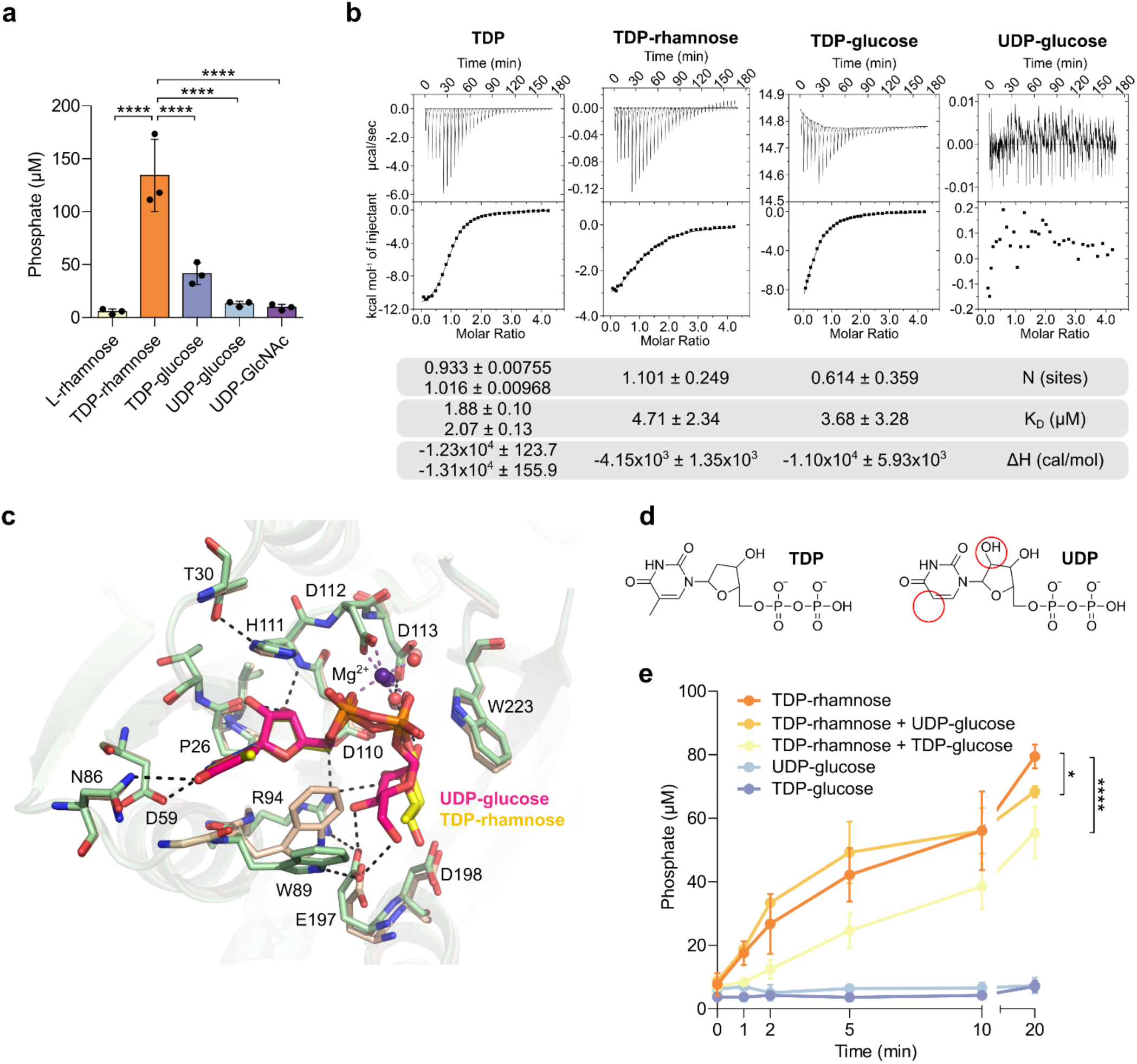
Donor substrate selectivity in RmlT. **a)** Purified recombinant RmlT (2 µM) was incubated with 1mM WTAs and 200 µM TDP-rhamnose, L-rhamnose, TDP-glucose, UDP-glucose or UDP-GlcNAc. RmlT activity was measured after 30 minutes by quantifying released phosphate. **b)** ITC measurement of RmlT/donor substrate interactions. Upper panels show the titration of RmlT with TDP-rhamnose, TDP-glucose, TDP or UDP-glucose. The lower panel shows heat released when substrate binds to RmlT as a function of relative concentrations. Dilution heat was corrected. Solid lines correspond to the best fit of data using a bimolecular interaction model. Table shows N-value (commonly referred as the stoichiometry), binding constant K_D_ (μM) and reaction enthalpy (ΔH) extracted from model fitting. **c)** Superposition of active-sites of RmlT/UDP-glucose (green with UDP-glucose in pink, PDB code: 8BZ6; chain D) and RmlT/TDP-rhamnose (wheat with TDP-rhamnose in yellow, PDB code: 8BZ7; chain D) through residues 467-477 (Cα RMSD: 0.114 Å) with the substrate-interacting residues shown as sticks. Hydrogen bonds between RmlT residues and UDP-glucose are represented as black dashed lines. Water and Mg^2+^ are shown as red and purple spheres, respectively. **d)** Chemical structure of TDP (left) and UDP (right) with differences highlighted by red circles. **e)** Quantification of phosphate released over time for RmlT incubated with different donor substrates. Reactions contained 1 mM WTAs, 200 µM of each nucleotide and 2 µM RmlT. Mean ± SD (n = 3, except for the curve of TDP-glucose in which n = 4) are shown for a) and e); individual values shown in a); two-way ANOVA; * *p* < 0.05, **** *p* < 0.0001.

To evaluate in more detail the molecular basis of RmlT specificity, we used isothermal titration calorimetry (ITC) to determine its affinity for different donor substrates in the absence of WTAs (Figure 5B). TDP-rhamnose bound to RmlT with a K_D_ of ∼13 µM, with 1 molecule bound per RmlT subunit. Surprisingly, TDP-glucose bound RmlT as well as TDP-rhamnose (K_D_ = ∼10 µM). Moreover, TDP also bound RmlT with a slightly lower affinity (K_D_ = ∼50 µM). We could not detect binding of UDP-glucose in the experimental conditions used. These results indicate that the nucleotide moiety (TDP) determines most of the strength of the interaction between the donor substrate and RmlT.

Using a high concentration (5 mM) of UDP-glucose, we determined the structure of its complex with RmlT (PDB code: 8BZ6) at 2.4 Å (Figure 5C, Table 1 and supplementary Table 1). The structure reveals few changes relative to RmlT/TDP-rhamnose and does not provide obvious hints for the preference towards TDP. The two nucleotides, TDP and UDP, differ in two chemical groups, a methyl group in the thymidine and a hydroxyl group in the ribose of UDP (Figure 5D). The methyl group in the base is within Van der Waals distance of Y28 but generally points away from the protein, while the extra hydroxyl in UDP is within hydrogen bonding distance of P26 main-chain carbonyl and appears to be accommodated with very slight changes in the protein. We also searched for clues that could explain a preference for rhamnose over glucose. Interestingly, the position and conformation of the sugar in UDP-glucose is significantly better defined than that of the rhamnose in the TDP-rhamnose structures (Supplementary Figures 3E-F, 5C and 9). This could reflect that the glucose is better stabilized in the sugar binding pocket resulting in reduced ability to acquire the proper position for catalysis when the acceptor WTAs is present. Moreover, while the glucose moiety in UDP-glucose establishes hydrogen bonds with E197 (Figure 5C), rhamnose in TDP-rhamnose is hydrogen bound to D198 (as discussed above).

Altogether, our results show that selection of the donor nucleotide does not result from interactions between the sugar and RmlT, but is determined in the ternary complex.

To gain insights into the donor substrate selectivity mechanism, we mixed RmlT with naked WTAs in the presence of the same concentrations of either TDP-rhamnose, TDP-glucose or UDP-glucose, and quantified RmlT activity by measuring released phosphate over time. As before, activity detected with TDP-rhamnose was much higher than with the other nucleotides (Figure 5E). We then measured RmlT activity in a competition experiment where the reaction contained equal amounts of TDP-rhamnose and TDP-glucose. If donor specificity is thermodynamically determined in the ternary complex, then the rate of the reaction should not change much with the addition of TDP-glucose since most of the ternary complexes will contain TDP-rhamnose. However, if the two donor nucleotides bind equally well and specificity is determined by another mechanism, we should observe a clear reduction in the rate of reaction, since only the fraction of the ternary complexes with bound TDP-rhamnose will result in sugar transfer. The time course of the TDP-rhamnose/TDP-glucose mixture showed a clear decrease in the initial activity rate calculated from the first 2 minutes, from 4.8 to 1.4 μM phosphate released/min/µM RmlT (Figure 5E). Moreover, when RmlT was incubated with mixed TDP-rhamnose and UDP-glucose (which binds very weakly to RmlT), the reaction occurred as in the presence of only TDP-rhamnose. In summary, these results support the proposal that RmlT specificity is not determined by the affinity of the donor substrate. Instead, the presence of rhamnose in the catalytic site increases either the binding affinity of WTAs or the efficiency of sugar transfer.

### RmlT rhamnosylates naked WTAs

We also wondered if RmlT displays specificity towards WTAs modifications, in particular WTAs glycosylations. We thus analysed RmlT activity in the presence of TDP-rhamnose and WTAs purified from wild-type EGDe strain and isogenic mutants presenting different glycosylation patterns: decorated with rhamnose at position *C*-4 and GlcNAc at *C*-2 of the ribitol-phosphate (RboP) units (isolated from wild-type *L. monocytogenes* EGDe strain), decorated only with GlcNAc (from the EGDe Δ*rmlT* strain), decorated only with rhamnose (from the EGDe Δ*lmo1079* strain), or without any decoration (naked WTAs) isolated from the EGDe Δ*rmlTΔlmo1079* strain. While some RmlT activity was detected with rhamnose-decorated WTAs, suggesting incomplete glycosylation at *C*-4, the strongest activity of RmlT was detected in the presence of naked WTAs (Figure 6A). Importantly, no activity was observed with WTAs containing only GlcNAc, indicating that the presence of GlcNAc impairs RmlT activity and strongly suggesting that in the bacterial cell, RmlT likely functions before the glycosyltransferase (Lmo1079) that adds GlcNAc on WTAs.

**Figure 6:**
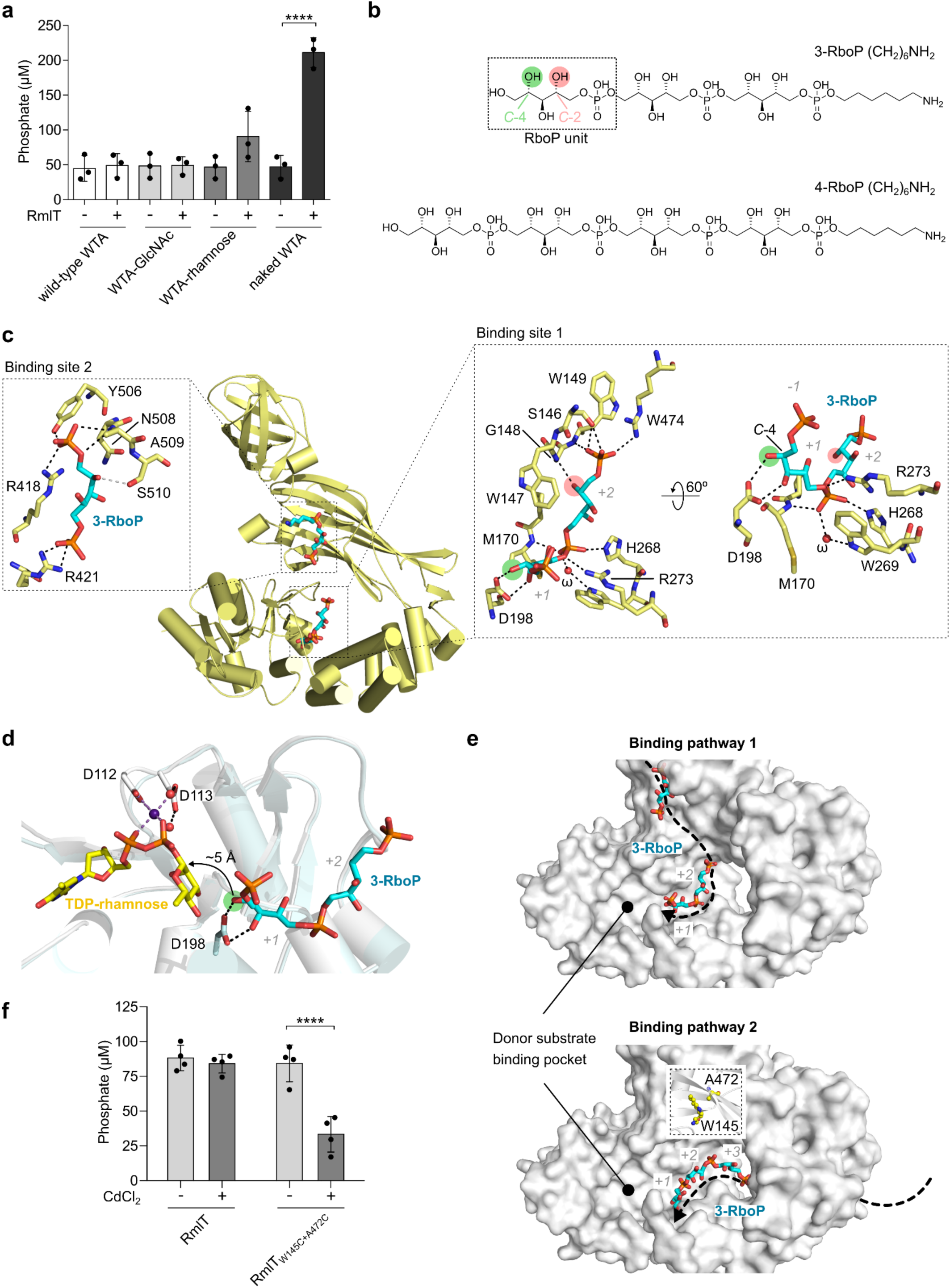
Structure of RmlT with bound poly-ribitol phosphate. **a)** Quantification of phosphate released from TDP generated by RmlT in presence of differently decorated WTAs: containing both GlcNAc and rhamnose modifications (wild-type WTAs), only GlcNAc modification (WTAs-GlcNAc), only rhamnose modification (WTAs-rhamnose) or without any glycosylation (naked WTAs). Reaction mixtures included 5 mM WTAs, 1 mM TDP-rhamnose and 10 μM RmlT incubated during 30 minutes. Mean ± SD (n = 3) and individual measurements are shown; one-way ANOVA, **** = p < 0.0001. **b)** Chemical structures of 3-RboP and 4-RboP with ribitol-phosphate unit indicated together with *C*-2 (light red) and *C*-4 (light green). **c)** Cartoon representation of RmlT with two bound RboP chains represented as sticks (PDB code 9GZJ chain A). Molecular details of RboP binding sites are shown, with subsites occupied by separate RboP units numbered +1 and +2. Atoms within hydrogen bond distance indicated by dashed lines. *C*-4 hydroxyl in subsite +1 is highlighted by green circle; *C*-2 hydroxyl in subsite +2 is highlighted by red circle. **d)** Comparison of active-sites in RmlT/TDP-rhamnose (white with TDP-rhamnose in yellow; PDB code 8BZ7 chain D) and RmlT/3-RboP (light cyan with 3-RboP in cyan; PDB code 9GZJ chain A). Structures are shown in same view after superposition through Cα atom of resides 238-516 (RMSD=0.364 Å). TDP/rhamnose, 3-RboP, residues D112 and D113 and D198 are shown as sticks. Mg^2+^ and water molecules are represented by purple and red spheres. Dashed lines indicate atoms within hydrogen bond or coordinating the metal. **e)** TarP/6RboP-(CH2)6NH2 (top image; PDB code 6HNQ chain A) and RmlT/3-RboP (bottom; PDB code 9GZJ chain A) in the same orientation (superposed through Cα atom pairs of RmlT residues 93-100 and 198-204 and TarP residues 75-82 and 181-187 (RMSD 0.672 Å)). Subsites are indicated (+1, +2 and +3). **f)** Quantification of phosphate released from TDP generated by RmlT or RmlT_W145C+A472C_ incubated with naked WTAs and TDP-rhamnose for 30 minutes in the presence or absence of 100 μM of CdCl_2_. Mean ± SD (n = 4) and individual measurement are shown; two-way ANOVA; **** p < 0.0001.

To better understand the molecular characteristics of the complex formed by RmlT and WTAs, we determined the structure of RmlT with two different synthetic WTAs, with 3 or 4 RboP units (Figure 6B), at 2.34 (PDB code: 9GZJ) and 2.52 Å (PDB code: 9GZK), respectively (Table 1 and supplementary Table 1). We also attempted to obtain diffracting crystals of the ternary complexes of RmlT with synthetic WTAs and either TDP-rhamnose or TDP-glucose, but diffraction quality was very poor (10 Å or worse). The RmlT/WTAs structures are globally similar to the apo-structure, with both structures showing detailed electron-density in all copies for 2 units of RboP in the vicinity of the catalytic residue D198 (binding site 1) (Figure 6C, Supplementary Figures 10 and 11). Interactions formed by the RboP units with the protein are detailed in Figure 6C. In particular, the phosphate group from the RboP in the subsite closest to the catalytic residues (subsite +1) interacts directly with side-chains of H268 and R273 (residues conserved in TarS, H253 and R258) and indirectly with W269 through a water molecule, as well as with the main-chain amino group of M170. In addition, the catalytic residue D198 is within hydrogen bond distance (2.8 and 2.9 Å) of the hydroxyls in *C*-3 and *C*-4, appearing to orient the *C*-4 hydroxyl group for nucleophilic attack on the sugar anomeric carbon in the donor substrate, as observed in a superposition of the binary complex structures with TDP-rhamnose and WTAs (Figure 6D). In addition, the RboP in subsite +2 (further from the catalytic site) establishes interactions through the phosphate group with S146, W149 and R474, through the hydroxyl in *C*-2 with the main-chain amino group of G148 and through Van der Walls contacts of *C*-1 and *C*-2 with the face of W147. Importantly, the WTAs structures provide an explanation for why RmlT prefers naked WTAs and cannot rhamnosylate WTAs modified with GlcNAc at *C*-2 since there is relatively little space around the hydroxyl at position *C*-2 in subsites +1 and +2 (Figure 6C).

Analysis of several structures provides clues for two possible interaction pathways taken by longer WTAs bound to RmlT. In the RmlT/WTAs structures, electron-density compatible with a second molecule of synthetic WTAs is present in some RmlT copies on the oligomerization domain (Figure 6C, Supplementary Figures 10 and 11). The lack of this density in other copies in the asymmetric unit seems linked to the formation of a crystal contact. We built a short WTAs into this density. Additionally, in the high resolution apo-RmlT structure (PDB code 8BZ5), HEPES molecules are present at different sites on the positively charged surface of the protein, with their sulphate groups nicely superposing with either the phosphate groups of RboP in binding sites 1 and 2 or with a strong electron-density observed in the RmlT/WTAs complex (not modelled) in binding site 3 (Supplementary Figure 12).

Altogether, this evidence suggests that, in binding pathway 1, long WTAs snakes through the oligomerization domain from binding site 2 into binding site 1, which is placed in a deep cleft close to the catalytic site (Figure 6E, Supplementary Figure 12).

However, a comparison of the RmlT/WTAs complex structures with the structures of TarP bound to similar synthetic WTAs (PDB codes: 6H4M, 6HNQ and 6H4F) (Figure 6E) suggests an alternative pathway. The comparison shows the WTAs sitting in equivalent clefts (corresponding to binding site 1) in the two structures (Figure 6E). The WTAs in the TarP structure have an extra RboP unit, corresponding to subsite +3, which in RmlT is positioned inside the tunnel formed by the catalytic, helical and oligomerization domains. This observation suggests that, in binding pathway 2, the long WTAs snakes through the tunnel into binding site 1, towards the catalytic site.

Binding pathway 2 raises the question of how would a long WTAs enter the tunnel. The strong structural similarity between RmlT and TarS (Supplementary Figure 1) [27] leads to the hypothesis that, like in TarS, the oligomerization domain of RmlT can adopt different positions relative to the catalytic domain so that there is a closed conformation, corresponding to the crystallographic structure, and an open conformation that allows access of the WTAs polymer to its binding groove and the tunnel. To explore this hypothesis, we generated a metal bridge across the catalytic/oligomerization interface by mutating two residues that are in close proximity to cysteine (W145 on the catalytic subdomain and A472 on the oligomerization subdomain) (Figure 6E) and performed the rhamnosylation reaction in the absence and presence of cadmium. Cadmium coordination by the two cysteines will bridge the two subdomains and stabilize the interface [31]. If a transition between the closed and open conformations is important for the rhamnosylation reaction of WTAs, then formation of the metal bridge will reduce activity. The mutant protein functioned as well as wild-type RmlT in the absence of cadmium cation, and the addition of 100 µM of CdCl_2_ had no impact the activity of wild-type RmlT. However, Cd^2+^ reduced the activity of the mutant protein by more than 50% (Figure 6F), supporting the proposal of the existence of an open conformation in the catalytic mechanism of RmlT and supporting WTAs binding pathway 2 as a viable option that merits future exploration.

### RmlT is a potential target for anti-virulence approaches

Previously, we demonstrated that deletion of the gene coding for RmlT in *L. monocytogenes* EGDe results in virulence attenuation in mouse spleens and livers, and that this virulence defect is reverted to wild-type levels by complementation with the wild-type RmlT (EGDe Δ*rmlT::rmlT*) [13, 14, 16, 17]. To further demonstrate the direct importance of the catalytic activity of RmlT for the virulence of *L. monocytogenes*, we assessed in the same mouse infection model the virulence of the *L. monocytogenes* EGDe Δ*rmlT* strain complemented with the catalytic-deficient RmlT_D198A_ mutant (EGDe Δ*rmlT::rmlT_D198A_*) and compared it with the Δ*rmlT* strain complemented with the wild type RmlT. We first demonstrated that similar protein levels of RmlT and RmlT_D198A_ are produced by wild-type *L. monocytogenes* EGDe and complemented EGDe Δ*rmlT* strains (Supplementary Figure 14). Mice were then intravenously infected with *L. monocytogenes* EGDe Δ*rmlT*, Δ*rmlT::rmlT* or Δ*rmlT::rmlT_D198A_*, and bacterial load per spleen and liver was assessed 72 h post-infection (Figure 7). Compared to the number of bacteria present in organs of animals infected with the *L. monocytogenes* EGDe Δ*rmlT* strain, the complementation with wild-type RmlT (EGDe Δ*rmlT::rmlT*) induced a significant increase in the number of bacteria present in both organs, restoring full bacterial virulence as previously observed [13]. In contrast, complementation with the inactive RmlT variant (EGDe Δ*rmlT::rmlT_D198A_*) did not increase the number of bacteria present in the organs of infected mice. These results indicate that inactivation of RmlT by a single mutation in its active site generates attenuated bacteria, serving as a proof of concept to demonstrate the potential of the inhibition of the catalytic activity of RmlT as a new anti-virulence approach.

**Figure 7:**
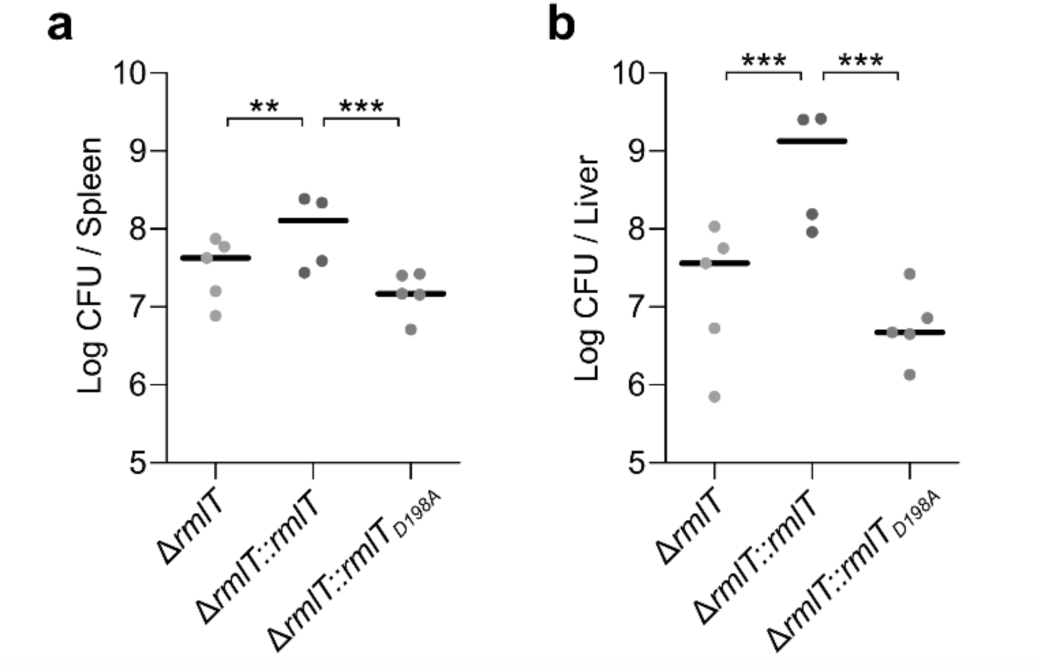
Impact of RmlT activity in the ability of *L. monocytogenes* to infect mice. Bacterial counts in the spleens (**a**) and livers (**b**) of the mice 72h after intravenous inoculation of the *L. monocytogenes* strains EGDe Δ*rmlT*, EGDe Δ*rmlT*::*rmlT* or EGDe Δ*rmlT*::*rmlT_D198A_*. Data are presented as scatter plots, with the counts for each animal indicated by dots and the mean by a horizontal line (n = 5, except for EGDe Δ*rmlT*::*rmlT* in which n = 4); one-way ANOVA; ** *p* < 0.01, *** *p* < 0.001.

## Discussion

Glycosylation of WTAs from Gram-positive pathogens is crucial for bacterial virulence and resistance to antimicrobials. Despite the significant impact of WTAs glycosyltransferases on crucial aspects of bacterial infection, identification and characterization of these enzymes, which represent promising targets for novel anti-virulence strategies, has been limited. In this study, we have elucidated the first structure and uncovered functional details of a bacterial WTAs rhamnosyltransferase, RmlT, expressed by the major Gram-positive foodborne pathogen *L. monocytogenes*. Structural analysis of RmlT reveals strong similarities with the structure of *S. aureus* glycosyltransferase TarS [27], both sharing the same tertiary organization in the catalytic, helical, oligomerization and turret subdomains. The catalytic and helical subdomains of these proteins are also identical in a third glycosyltransferase, *S. aureus* TarP [24]. In addition, many of the residues present in the active site of RmlT are conserved in TarS and TarP [24, 27]. The most striking difference is that RmlT assembles as a dimer while TarS and TarP form trimers [24, 27]. We showed that the dimer interface is important for the physiological function of RmlT since mutations on the interface designed to disrupt the dimer architecture retained catalytic activity but altered the rhamnosylation pattern of the bacterial surface *in vivo*.

Although we have previously demonstrated that in *L. monocytogenes* the gene coding for RmlT is required for the decoration of WTAs with rhamnose [13], the activity of the enzyme had never been characterized. We now have shown, using two different assays, that RmlT in the presence of TDP-rhamnose and naked WTAs alters the electrophoretic mobility of WTAs in a native-PAGE and releases TDP, consistent with the transfer of rhamnose from the donor substrate to WTAs. Given the high conservation of the catalytic residues across different WTAs glycosyltransferases, we wondered what drives the specificity of RmlT activity for TDP-rhamnose. Using a combination of approaches, we showed that RmlT binds equally well to TDP-rhamnose and TDP-glucose but only transfers rhamnose to WTAs, suggesting that specificity is either kinetically determined or depends on a cooperative binding effect between the WTAs and the correct donor substrate. Similar results were reported for *B. subtilis* TagA, a teichoic acid glycosyltransferase. TagA binds both UDP-*N*-acetylmannosamine (UDP-ManNAc) and UDP-GlcNAc but is enzymatically active only with UDP-ManNAc [32]. This finding raises the question of how the efficiency of WTAs rhamnosylation is ensured in bacteria since other TDP coupled sugars will compete for the TDP-rhamnose binding site, slowing the production of decorated WTAs.

Our structures provide a clarification for the role of the catalytic residue D198, which aligns with the proposed catalytic residue in *S. aureus* Tar proteins [24, 27]. In the binary complex with the acceptor substrate, D198 forms hydrogen bonds with *C*-3 and *C*-4 of the ribitol-phosphate at the catalytic site, suggesting that D198 catalyses the reaction by orienting the hydroxyl towards the anomeric carbon of the sugar and by acting as a base that promotes the nucleophilic character of the hydroxyl group.

In addition, we showed that RmlT displays specificity for WTAs, glycosylating naked WTAs that lacks both rhamnose and GlcNAc. The binary complex structure with synthetic WTAs reveals that this RmlT preference results from steric hindrance at *C*-2 substitutions.

Overall, our data provides a novel insight into the molecular mechanism of WTAs glycosyltransferases. The RmlT structure in complex with WTAs offers clues about the long WTAs pathway on the protein surface. In particular, a second RboP binding site suggests that the polymer runs along the oligomerization domain into the deep cleft that leads to the catalytic site. Alternatively, comparison between the structures of RmlT and TarP and biochemical crosslinking across the interface between the RmlT catalytic and oligomerization subdomains, suggests that RmlT undergoes a transition between an open conformation, where the catalytic-oligomerization interface is not established and the WTAs polymer can bind to the protein, and a closed conformation, where the interface is formed and the polymer is trapped inside the tunnel formed by the catalytic, oligomerization and helical subdomains. In this alternative pathway, long WTAs run through the tunnel towards the catalytic site. Further studies are required to distinguish between these two possibilities and their impact on the lifetime of the enzyme-WTAs complex and on the potential processivity of the enzyme as proposed for TarS and TarP (Figure 8) [24, 27].

**Figure 8:**
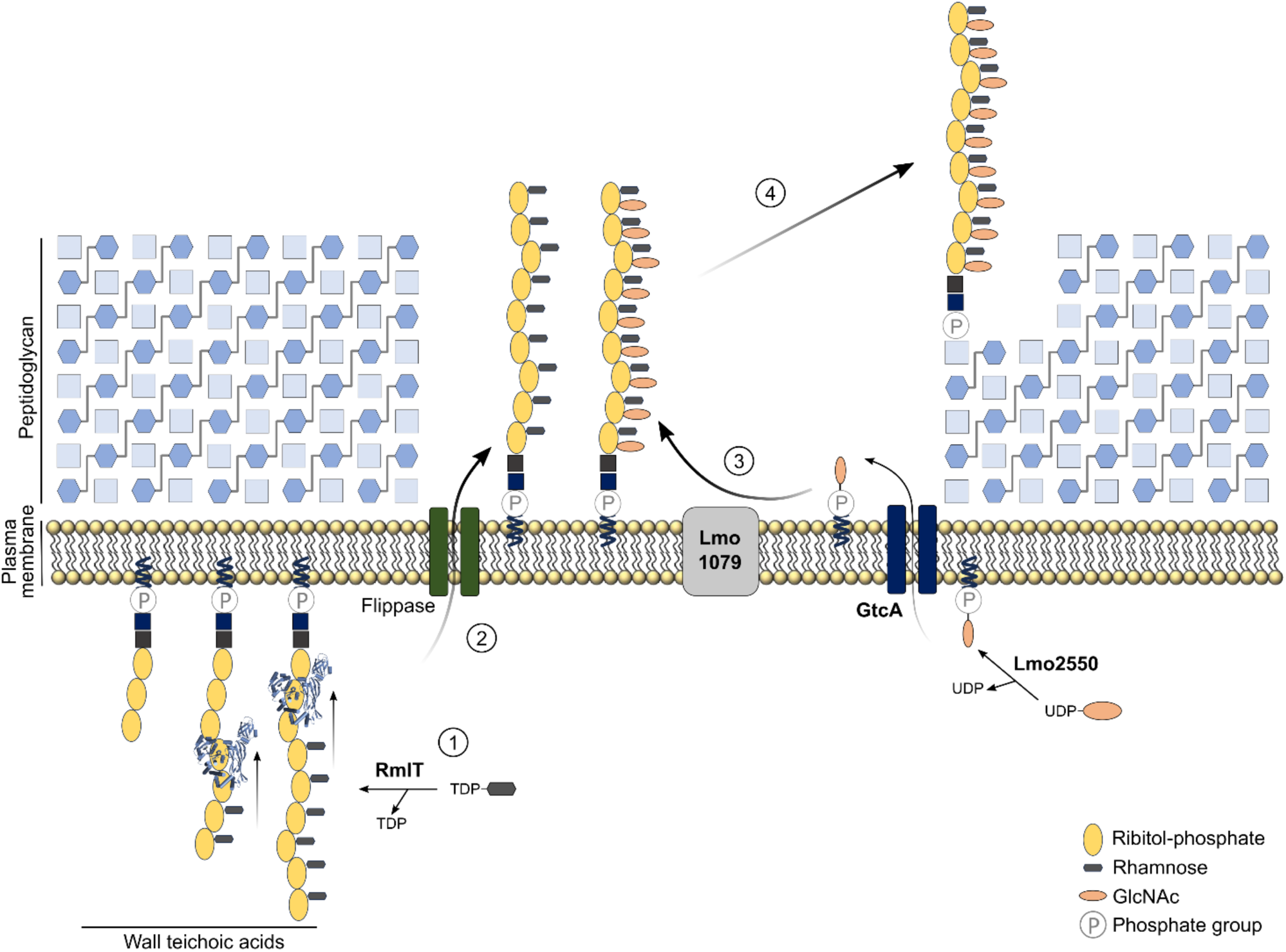
Proposed model for the glycosylation of WTAs in *L. monocytogenes* EGDe serovar 1/2a. Step 1: In the cytoplasm, RmlT decorates newly formed chains of ribitol-phosphate with rhamnose units from TDP-rhamnose. Step 2: Rhamnosylated WTAs are flipped across the membrane, probably by the Lmo1074-75 complex, homolog to the *S. aureus* two-component ATP-binding cassette (ABC) transporter TarGH. Step 3: Rhamnosylated WTAs are glycosylated by Lmo1079 with GlcNAc from a C_55_-P-GlcNAc intermediate previously formed by the cytoplasmic glycosyltransferase Lmo2550 using UDP-GlcNAc and then flipped across the membrane by the GtcA flippase. Step 4: Fully decorated WTAs are anchored to the cell wall, possibly by a homolog of the *S. aureus* TarTUV complex.

By combining our findings with earlier discoveries, we clarified several stages of WTAs glycosylation in *L. monocytogenes* serovar 1/2a. In particular, our data supports that the newly formed WTAs is first decorated with rhamnose by RmlT. This indicates that decoration with GlcNAc by Lmo1079 follows rhamnosylation, fitting well with the physiological location of RmlT in the cytosol and of Lm1079 on the external surface of the plasma membrane (Figure 8). We also suggest that rhamnosylation (step 1) involves a close interaction of RmlT with a long stretch of ribitol-phosphate units in the WTAs, favouring multiple cycles of rhamnosylation before release of the decorated WTAs. Flipping of the rhamnosylated WTAs to the outside of the cell membrane (step 2) is likely to result from the action of a homolog of the *S. aureus* TarGH ABC transporter (predicted to be Lmo1074-75) [35, 36]. In parallel, it has been proposed that the cytoplasmic glycosyltransferase Lmo2550 uses UDP-GlcNAc to form a C_55_-P-GlcNAc WTAs intermediate that is also flipped across the membrane by the flippase GtcA [29]. The membrane-associated glycosyltransferase Lmo1079 then transfers the GlcNAc moiety to the rhamnosylated-WTAs backbone on the outside of the cell [37, 38] (step 3). Finally, the mature WTAs is transferred to the bacterial cell wall by a still unclear mechanism proposed to be promoted in *S. aureus* by the TarTUV enzymes (step 4) [33].

In summary, WTAs glycosylations are involved in many bacterial features across different species including bacterial virulence [13, 14], resistance to antibiotics [16, 23, 25, 39–42] and antimicrobial peptides [13] and in the localization of important surface proteins that facilitate host immune evasion and colonization [23, 25, 39, 40]. Therefore, it is tempting to consider glycosylation of bacterial WTAs as a conserved virulence mechanism in Gram-positive pathogens. Importantly, our experiments proved that mutation of a single residue in the active site of RmlT leads to enzyme inactivation and to a significant decrease in the capacity of *L. monocytogenes* cells to infect mice. Together with the relatively weak binding selectivity for TDP-modified molecules, this indicates that RmlT is a valid target for the development of next-generation antibacterial drugs that, by inhibiting specific sugar decorations, will not directly kill bacteria but will diminish bacterial virulence, increase susceptibility to host innate defences and potentiate the action of antibiotics while preserving the microbiota.

## Methods

### Cloning

*rmlT* was cloned into the expression vector pET28b by the polymerase incomplete primer extension method [43]. Briefly, *rmlT* was amplified by PCR from the *L. monocytogenes* EGDe (ATCC-BAA-679) genomic DNA, using primers #1 and #2. Vector pET28b was amplified by PCR using primers #3 and #4. Both reactions generated a mixture of incomplete extensions products that contained short overlapping sequences at the ends, allowing the annealing of the complementary strands to produce hybrid vector-insert combinations. Vectors and primer sequences are presented in Supplementary Table 2 and 3, respectively.

### Site-directed mutagenesis

Vectors pET28b-*rmlT*_D198A,_ pET28b-*rmlT*_D197A,_ pET28b-*rmlT*_W89A,_ pET28b-*rmlT*_W223A,_ pET28b-*rmlT*_Y368A+E372A,_ pET28b-*rmlT*_Y347A+Y368A+E372A_, pET28b-*rmlT*_W145C+A472C_, pPL2-*rmlT*_D198A_, pPL2-*rmlT*_D197A_, pPL2-*rmlT*_W89A_, pPL2-*rmlT*_W223A_, pPL2-*rmlT*_Y368A+E372A_ and pPL2-*rmlT*_Y347A+Y368A+E372A_ were obtained by site-directed mutations using pET28b-*rmlT* or pPL2-*rmlT* [13] as templates and primer pairs #5 and #6 (for pET28b-*rmlT*_D198A_ and pPL2-*rmlT*_D198A_), #7 and #8 (for pET28b-*rmlT*_D197A_ and pPL2-*rmlT*_D197A_), #9 and #10 (for pET28b-*rmlT*_W89A_ and pPL2-*rmlT*_W89A_), #11 and #12 (for pET28b-*rmlT*_W223A_ and pPL2-*rmlT*_W223A_), #13 and #14 (for pET28b-*rmlT*_Y368A+E372A_ and pPL2-*rmlT*_Y368A+E372A_), #13, #14 #15 and #16 (for pET28b-*rmlT*_Y347A+Y368A+E372A_ and pPL2-*rmlT*_Y347A+Y368A+E372A_) and #17, #18, #19, #20 (for pET28b-*rmlT*_W145C+A472C_). Primer information is presented in Supplementary Table 3. Mutations were confirmed by sequencing.

### Synthesis of 3RboP and 4RboP

3RboP and 4RboP were synthesized as previously described [39, 44].

### Expression and purification of RmlT variants

*Escherichia coli* BL21(DE3) cells transformed with pET28b encoding N-terminally hexahistidine tagged RmlT (pET28b-*rmlT*), RmlT_D198A_ (pET28b-*rmlT*_D198A_), RmlT_D197A_ (pET28b-*rmlT*_D197A_), RmlT_W89A_ (pET28b-*rmlT*_W89A_), RmlT_W223A_ (pET28b-*rmlT*_W223A_), RmlT_Y368A+E372A_ (pET28b-*rmlT* _Y368A+E372A_), RmlT_Y347A+Y368A+E372A_ (pET28b-*rmlT* _Y347A+Y368A+E372A_) or RmlT_W145C+A472C_ (pET28b-*rmlT* _W145C+A472C_) were grown in LB supplemented with 30 μg ml^−1^ kanamycin at 37°C for 3 h. Protein expression was induced with 0.5 mM isopropyl β-D-1-thiogalactopyranoside (IPTG). After 16 h-growth at 30°C, cells were harvested, resuspended in lysis buffer (50 mM NaH_2_PO_4_ pH 8.0, 300 mM NaCl, 10 mM Imidazole) and lysed by sonication. Cell lysate was clarified by centrifugation and incubated with an immobilized metal-affinity matrix (Ni-NTA Agarose, ABT Technologies), pre-equilibrated with lysis buffer, at 4°C for 1 h. After washing the beads with 50 mM NaH_2_PO_4_ pH 8.0, 300 mM NaCl, 30-50 mM Imidazole, proteins were eluted with 50 mM NaH_2_PO_4_, 300 mM NaCl, pH 8.0, 100-300 mM Imidazole. Eluted material was concentrated using a 30 kDa cut-off membrane concentrator (Vivaspin, Sartorius Stedim Biotech) concomitantly with buffer exchange to 20 mM Tris-HCl pH 8.0, 100 mM NaCl, 1 mM DTT. Hexahistidine tag (His-tag) was removed by digestion with TEV protease (1:10 molar ratio) at 4°C for 16 h. Proteins were separated from TEV protease, His-tag and uncut proteins by a second immobilized metal-affinity column pre-equilibrated with storage buffer (20 mM Tris-HCl pH 8.0, 100 mM NaCl, 1 mM DTT). Flow-through containing untagged protein was concentrated and loaded into a Superdex-200 10/300 GL column (GE Healthcare) pre-equilibrated with storage buffer. Fractions containing pure protein were pooled and concentrated. Protein concentration was estimated by measuring absorbance at 280 nm and using extinction coefficients calculated with the ExPASY tool Protparam. Purified RmlT variants were flash-frozen in liquid nitrogen and stored at −80 °C.

### Expression and purification of selenomethionine-containing RmlT variant

*E. coli* B834(DE3) cells transformed with pET28b-*rmlT* were grown in 20 mL LB medium overnight at 37°C. Cells were collected, washed three times with sterile deionized water and inoculated into selenomethionine medium (Molecular Dimensions). Protein expression and purification was performed as described above.

### Crystallization of RmlT variants

Initial crystallization conditions for RmlT were determined at 20°C using commercial sparse-matrix crystallization screens. Sitting-drop vapor diffusion experiments were set up in 96-well CrystalQuick plates using an Oryx 4 crystallization robot (Douglas Instruments). Drops consisted of equal volumes (0.3 μL) of RmlT protein solution (at 10 mg mL^−1^ in 20 mM Tris-HCl pH 8.0, 100 mM NaCl, 1 mM DTT) and crystallization solution, equilibrated against 40 μL reservoir solution. The apo RmlT structure with bound HEPES was obtained from a crystal grown in solution B9 from JBScreen Wizard 1 & 2 crystallization screen [100 mM HEPES-NaOH pH 7.5, 20% (w/v) PEG 8,000]. Crystal was frozen directly in liquid nitrogen. Selenomethionine-containing crystals (apo structure) were first obtained in solution H5 from NeXtal Cystal Classic II Suite crystallization screen (100 mM succinic acid pH 7.0, 15% (w/v) PEG 3,350). Crystals were reproduced using MRC Maxi 48 well plates with drops composed of equal volumes (1 μL) of protein (at 10 mg mL^−1^ in 20 mM Tris-HCl pH 8.0, 100 mM NaCl, 1 mM DTT) and crystallization solution (200 mM succinic acid pH 7.0, 10% (w/v) PEG 3,350) equilibrated against 150 μL reservoir. Crystals were cryo-protected with a 1:1 mixture of Paratone N and Mineral oil before flash freezing in liquid nitrogen.

The RmlT/TDP-rhamnose (Thymidine-5’-diphosphate-L-rhamnose disodium, Biosynth Ltd, UK) structure was determined from soaking a crystal obtained in 100 mM Bis-Tris Propane pH 7.0, 200 mM K/Na tartrate tetrahydrate, 20% (w/v) PEG 3,350 (originally identified in condition F9 from PACT premier (Molecular Dimensions). Before freezing, 0.5 μL of a solution containing 55 mM Bis-Tris propane pH 7.0, 110 mM K/Na tartrate tetrahydrate, 13.75% (w/v) PEG 3,350, 5% (v/v) glycerol, 25 mM MgCl_2_ and 30 mM TDP-rhamnose was added to the crystal drop and allowed to equilibrate for 50 min. After quickly transferring the crystal into a solution containing 220 mM K/Na tartrate tetrahydrate, 110 mM Bis-Tris propane pH 7.0, 27.5% (w/v) PEG 3,350, 10% (v/v) glycerol, the crystal was flash-frozen in liquid nitrogen. The structure of RmlT_D198A_ in complex with TDP-rhamnose was determined from a crystal grown in 100 mM Bis-Tris propane pH 7.0, 200 mM K/Na tartrate, 22.5% (w/v) PEG 3,350. Crystal was transferred to a 0.5 μL of reservoir solution and then 0.5 μL of 55 mM Bis-Tris propane pH 7.0, 110 mM K/Na tartrate tetrahydrate, 16.75% (w/v) PEG 3,350, 20 mM MgCl_2_, 20 mM TDP-rhamnose was added. The mixture was allowed to equilibrate for 6 min. Cryoprotection was achieved by adding 0.8 μL of a solution containing 110 mM Bis-Tris propane pH 7.0, 220 mM K/Na tartrate tetrahydrate, 33.5% (w/v) PEG 3,350 prior to flash-freezing in liquid nitrogen. Finally, RmlT crystals were also obtained in condition H6 of the NeXtal Classics Lite Suite crystallization screen (100 mM Tris-HCl pH 8.5, 200 mM sodium acetate., 15% (w/v) PEG 4,000). A crystal grown in this condition was used to solve the RmlT/UDP-glucose structure. Crystal was sequentially transferred to solutions with increasing concentration of DMSO and supplemented with UDP-glucose (110 mM Tris-HCl pH 8.5, 220 mM sodium acetate, 16.5% (w/v) PEG 4,000, 10 to 30% DMSO, 5 mM MgCl2, 5 mM UDP-glucose) before flash-freezing in liquid nitrogen.

The <u>RmlT structure with bound synthetic 3RboP-(CH_2_)_6_NH_2_</u> was obtained from a crystal grown with equal volumes (1 μL) of RmlT protein solution (at 10 mg mL^−1^ in 20 mM Tris-HCl pH 8.0, 100 mM NaCl, 1 mM DTT, 5 mM MgCl_2_ and with 4 mM 3RboP-(CH_2_)_6_NH_2_) and crystallization solution (100 mM MES pH 6.5, 150 mM K/Na tartrate tetrahydrate, 20% (w/v) PEG 3,350) equilibrated against 40 μL reservoir solution. The crystal was transferred to 0.5 μL of crystallization solution and 0.5 uL was added of a solution containing 99 mM MES pH 6.5, 148.5 mM K/Na tartrate tetrahydrate, 19.8% (w/v) PEG 3,350, 18% glycerol, 5 mM 3RboP- (CH_2_)_6_NH_2_ and 5 mM MgCl_2_ before flash freezing in liquid nitrogen. The RmlT structure with bound synthetic 4RboP-(CH_2_)_6_NH_2_ was obtained from a crystal grown with equal volumes (1 μL) of RmlT protein solution (at 10 mg mL^−1^ in 20 mM Tris-HCl pH 8.0, 100 mM NaCl, 1 mM DTT, 5 mM MgCl_2_ and with 4 mM 4RboP-(CH_2_)_6_NH_2_) and crystallization solution (100 mM Bis-Tris propane pH7.0, 200 mM K/Na tartrate tetrahydrate, 17.5% (w/v) PEG 3,350) equilibrated against 40 μL reservoir solution. The crystal was cryo-protected with a 1:1 mixture of Paratone N and Mineral oil before flash-freezing in liquid nitrogen.

### Data collection and processing

Diffraction data were collected from cryo-cooled (100 K) single crystals with a ∼0.98 Å wavelength X-ray beam at beamlines PROXIMA-1 and PROXIMA-2A of the French National Synchrotron Facility (SOLEIL, Gif-sur-Yvette, France) and XALOC of ALBA (Barcelona, Spain). Data sets were processed with XDS [45] and reduced with utilities from the CCP4 program suite [46]. X-ray diffraction data statistics are summarized in Table 1.

### Structure determination, model building and refinement

The structure of apo RmlT was solved by single-wavelength anomalous diffraction (SAD), using the anomalous signal of the selenium at the *K*-absorption edge and pipelines from CCP4 package [46]: the SHELXC/SHELXD/SHELXE pipeline [47] was used for Se atom location, the Phaser SAD pipeline [48, 49] for density modification and automatic model building and the Buccaneer pipeline [50, 51] used for automatic model building and refinement. All the other structures were solved by molecular replacement with Phaser [48] using chain A of the apo structure as a search model. Alternating cycles of model building with Coot [52] and refinement with PHENIX [53] were performed until model completion. Simulated annealing was performed once for structures solved by molecular replacement. NCS restrains were used only in the first cycles of refinement. TLS refinement was performed in more advanced stages of refinement, with TLS groups identified automatically by PHENIX. Refined coordinates and structure factors were deposited at the Protein Data Bank [54]. The PDB accession codes 8BZ4, 8BZ5, 8BZ6, 8BZ7, 8BZ8, 9GZJ and 9GZK were assigned to apo RmlT, apo RmlT with HEPES, RmlT in complex with UDP-glucose, RmlT in complex with TDP-rhamnose, and RmlT_D198A_ in complex with TDP-rhamnose, RmlT in complex with 3-RboP-(CH2)6NH2 and RmlT in complex with 4-RboP-(CH_2_)_6_NH_2_, respectively. Refinement statistics are summarized in Table 1.

### Analysis of crystallographic structures

The crystallographic models were superposed with SUPERPOSE [46, 55] and secondary structure elements were identified with PROMOTIF [56]. The surface electrostatic potential was calculated with APBS [57] using the AMBER force field [58]. Figures depicting molecular models were created with PyMOL (Schrӧdinger).

### Isothermal titration calorimetry

Recombinant RmlT was dialysed overnight against 20 mM HEPES pH 8.0, 100 mM NaCl, 5 mM MgCl_2_. Solution of 200 µM TDP, TDP-rhamnose, TDP-glucose or UDP-glucose (calorimetric syringe) were titrated into 20 µM RmlT (sample cell), with an initial injection of 2 µL, followed by 28 injections of 10 µL. Titrations were performed in a MicroCal VP-ITC instrument (GE Healthcare) at 15°C. Data were visualized and analyzed with Origin 7 Software using a single-site binding model.

### Analytical size-exclusion chromatography

Analytical size-exclusion chromatography was performed on a Superdex 200 10/300 GL column (GE Healthcare) pre-equilibrated with protein storage buffer. Blue dextran (2,000 kDa), Ferritin (440 kDa), Aldolase (158 kDa), Ovalbumin (43 kDa) and Ribonuclease A (13.7 kDa) were used as standards for column calibration. The K_av_ *versus* Log molecular weight was calculated using the equation *K_av_* = (*V_e_* − *V_0_*)⁄(*V_t_* − *V_0_*), where *V_e_* is the elution volume of the protein, *V_0_* is the void volume of the column and *V_t_* is the column bed volume.

### Listeria monocytogenes strains

We previously generated the *L. monocytogenes* strains EGDe Δ*rmlT*, EGDe Δ*rmlT*::*rmlT* [13] and EGDe Δ*rmlT*Δl*mo1079* [16]. To generate the *L. monocytogenes* strains EGDe Δ*rmlT*::*rmlT*_D198A_, EGDe Δ*rmlT*::*rmlT*_D197A_, EGDe Δ*rmlT*::*rmlT*_W89A_, EGDe Δ*rmlT*::*rmlT*_W223A_, EGDe Δ*rmlT*::*rmlT*_Y368A+E372A_ and EGDe Δ*rmlT*::*rmlT*_Y347A+Y368A+E372A_ strains, the pPL2-derivative plasmids described above were introduced into *E. coli* S17–1 and transferred to the *L. monocytogenes* EGDe Δ*rmlT* strain by conjugation on Brain Heart Infusion (BHI) agar. Transconjugant clones were selected in BHI supplemented with chloramphenicol/ colistin/nalidixic acid (Cm/Col/Nax) and chromosomal integration of the plasmids confirmed by PCR and sequencing.

### Quantitative proteomics analysis

To quantify the level of RmlT in the different *L. monocytogenes* mutants, cytosolic proteomes were isolated. Bacteria cultures growing in BHI were harvested at late exponential growth phase (OD_600nm_ ≈ 0,8) by centrifugation (4 000 xg for 10 min) and bacterial cells washed twice in lysis buffer (50 mM Tris-HCl (pH 7.5), 1 % SDS and Protease Inhibitor Cocktail (cOmplete™, Roche). Bacteria were mechanically lysed by bead beating in a FastPrep apparatus (45 s, speed 6.5 three cycles) and centrifuged at 20 000 xg for 30 min to remove cellular debris. Supernatant containing the cytosolic soluble proteins was recovered and total protein was quantified using the Bradford method. Each sample was processed for proteomic analysis following the solid-phase-enhanced sample-preparation (SP3) protocol and enzymatically digested with Trypsin/LysC as previously described [59]. Protein identification and quantitation was performed by nanoLC-MS/MS equipped with a Field Asymmetric Ion Mobility Spectrometry - FAIMS interface. This equipment is composed of a Vanquish Neo liquid chromatography system coupled to an Eclipse Tribrid Quadrupole, Orbitrap, Ion Trap mass spectrometer (Thermo Scientific, San Jose, CA). 250 nanograms of peptides of each sample were loaded onto a trapping cartridge (PepMap Neo C18, 300 μm x 5 mm i.d., 174500, Thermo Scientific, Bremen, Germany). Next, the trap column was switched in-line to a μPAC Neo 50 cm column (COL-nano050NeoB) coupled to an EASY-Spray nano flow emitter with 10 μm i.d. (ES993, Thermo Scientific, Bremen, Germany). A 130 min separation was achieved by mixing A: 0.1% FA and B: 80% ACN, 0.1% FA with the following gradient at a flow of 750 nL/min: 0.1 min (1% B to 4% B) and 1.9 min (4% B to 7% B). Next, the flow was reduced to 250 nL/min with the following gradient: 0.1 min (7.0 to 7.1% B), 80 min (7.1% B to 22.5% B), 30 min (22.5% B to 40% B), 8 min (40%B to 99% B) and 9.9 min at 99% B. Subsequently, the column was equilibrated with 1% B. Data acquisition was controlled by Xcalibur 4.6 and Tune 4.0.4091 software (Thermo Scientific, Bremen, Germany). MS results were obtained following a Data Dependent Acquisition - DDA procedure. MS acquisition was performed with the Orbitrap detector at 120 000 resolution in positive mode, quadrupole isolation, scan range (m/z) 375-1500, RF Lens 30%, standard AGC target, maximum injection time was set to auto, 1 microscan, data type profile and without source fragmentation. FAIMS mode: standard resolution, total carrier gas flow: static 4L/min, FAIMS CV: −45, −60 and −75 (cycle time, 1 s).

Internal Mass calibration: Run-Start Easy-IC. Filters: MIPS, monoisotopic peak determination: peptide, charge state: 2-7, dynamic exclusion 30s, intensity threshold, 5.0e3. MS/MS data acquisition parameters: quadrupole isolation window 1.8 (m/z), activation type: HCD (30% CE), detector: ion trap, IT scan rate: rapid, mass range: normal, scan range mode: auto, normalized AGC target 100%, maximum injection time: 35 ms, data type centroid. The raw data were processed using the Proteome Discoverer 3.1.0.638 software (Thermo Scientific) and searched against the UniProt database for the *Listeria monocytogenes* EGDe Proteome (2024_02 with 2,844 entries). A common protein contaminant list from MaxQuant was also included in the analysis. The Sequest HT search engine was used to identify tryptic peptides. The ion mass tolerance was 10 ppm for precursor ions and 0.5 Da for fragment ions. The maximum allowed missing cleavage sites was set to two. Cysteine carbamidomethylation was defined as constant modification. Methionine oxidation, deamidation of glutamine and asparagine, peptide terminus glutamine to pyroglutamate, and protein N-terminus acetylation, Met-loss, and Met-loss+acetyl were defined as variable modifications. Peptide confidence was set to high. The processing node Percolator was enabled with the following settings: maximum delta Cn 0.05; target FDR (strict) was set to 0.01 and target FDR (relaxed) was set to 0.05, validation based on q-value. Protein label-free quantitation was performed with the Minora feature detector node at the processing step. Precursor ions quantification was performed at the consensus step with the following parameters: unique plus razor peptides were included, precursor abundance based on intensity, and normalization based on total peptide amount. For hypothesis testing, protein ratio calculation was pairwise ratio-based and an t-test (background based) hypothesis test was performed.

### Isolation and purification of wall teichoic acids

WTAs from *L. monocytogenes* strains wild-type, Δ*rmlT*, Δ*lmo1079* and Δ*rmlT*Δ*lmo1079*, were isolated and purified as previously described [60] with some modifications. *L. monocytogenes* cells grown overnight in 20 mL BHI at 37°C were collected by centrifugation at 4,000xg for 10 min. Cells were washed once in buffer 1 (50 mM MES pH 6.5), resuspended in buffer 2 (50 mM MES pH 6.5, 4% (w/v) SDS), and boiled in a water bath for 1 h. Cell debris was collected by centrifugation at 4,000xg for 10 min and resuspended in 2 mL of buffer 1. The pellet was washed sequentially in 1 ml of buffers 2, 3 [50 mM MES pH 6.5, 2% (w/v) NaCl] and 1. Washed pellet was resuspended in 1 mL of 20 mM Tris-HCl pH 8.0, 0.5% (w/v) SDS, 20 μg proteinase K and incubated at 50°C for 4 h with agitation (1,400 rpm). To remove SDS, the sample was centrifuged at 16,000xg for 1 min and washed once with buffer 3 and three times with distilled H_2_O. Sample was then resuspended in 1 mL of 0.1 M NaOH and shaken at room temperature for 16 h. The remaining insoluble cell debris was removed by centrifugation at 16,000xg for 10 min and the supernatant containing the hydrolysed crude WTAs was neutralized with 1 M Tris-HCl pH 7.8. The sample was dialyzed against distilled H_2_O using a 1 kDa cut-off membrane. Finally, isolated crude WTAs were lyophilized and resuspended in ultra-pure H_2_O to a final concentration of 10 mg mL^−1^ (considering ribitol monomer molecular weight) [60].

### RmlT activity assay

The activity of RmlT variants was analyzed using the Glycosyltransferase Activity Kit (R&D Systems) according to the manufacturer’s protocol. Recombinant RmlT (2 μM), was incubated with wild-type WTAs, containing both GlcNAc and rhamnose modifications (wild-type WTAs purified from the *L. monocytogenes* strains EGDe); WTAs containing only GlcNAc modification (WTAs-GlcNAc purified from EGDe Δ*rmlT*); WTAs containing only rhamnose modification (WTAs-rhamnose purified from EGDe Δ*lmo1079*); and naked WTAs without any glycosylation (naked WTAs purified from EGDe Δ*rmlT*Δ*lmo1079*) (1 mM) and TDP-rhamnose, UDP-rhamnose or UDP-glucose (200 μM) in reaction buffer (5 mM MgCl_2_, 25 mM Tris, 10 mM CaCl_2_, pH 7.5) at 37°C for 30 min. In the metal bridge coordination experiment with RmlT_W145C+A472C_, the enzyme (2 μM) was incubated with naked WTAs (1 mM) and TDP-rhamnose (200 μM) in reaction buffer (5 mM MgCl_2_, 25 mM Tris, 10 mM CaCl_2_, 100 μM TCEP, pH 7.5) at 37°C for 30 min, in presence of absence of CdCl_2_ (100 μM). Free phosphate released upon hydrolysis of TDP or UDP produced by RmlT was quantified by absorbance at 620 nm using a Synergy 2 multi-mode plate reader (BioTek, USA). RmlT reaction mixtures were also separated in 20 % polyacrylamide gels and WTAs were stained with 1 mg mL^−1^ Alcian Blue solution after overnight incubation.

### Cell wall rhamnose binding assay

Rhamnose present at the bacterial surface was quantified using a bacteriophage receptor binding protein (gp17) fused to GFP (gp17-GFP) (provided by M. Loessner, ETH Zurich). The assay was performed as described before [18]. Briefly, purified gp17-GFP was centrifuged at 30,000xg and 4°C for 1 h to remove protein aggregates. Cells from one milliliter of exponentially growing bacterial (OD_600nm_ = 0.5-0.6) were harvested by centrifugation, resuspended in 100 µL of PBS and incubated with 2.4 μg of gp17-GFP at room temperature for 20 min. Cells were washed three times with PBS and binding to the bacterial cell wall was visualized using a fluorescence microscope Olympus BX53 or quantified using a plate reader Synergy 2 (BioTek, USA).

### Animal infections

Virulence studies were performed in mouse models of BALB/c strain. Infections were performed in six-to-eight week-old specific-pathogen-free females (n=5, except for EGDe Δ*rmlT*::*rmlT* in which n=4) as described [61]. Briefly, intravenous infections were performed through the tail vein with 10^4^ CFU of bacteria in PBS. The infection was carried out for 72 h, at which point animals were euthanatized by general anaesthesia. Spleen and liver were aseptically collected, homogenized in sterile PBS, and serial dilutions of the organ homogenates plated in BHI agar. Mice were maintained at the i3S animal facilities, in high efficiency particulate air (HEPA) filter-bearing cages under 12 h light cycles, and were given sterile chow and autoclaved water *ad libitum*. All animal procedures were performed in agreement with the guidelines of the European Commission for the handling of laboratory animals (directive 2010/63/EU), the Portuguese legislation for the use of animals for scientific purposes (Decreto-Lei 113/2013), and were approved by the i3S Animal Ethics Committee as well as by the Direcção Geral de Veterinária (the Portuguese authority for animal protection) under licenses 2017-06-30 015302 and 2023-04-05 006564.

### Statistical analyses

Statistical analyses were calculated using the software Prism 9 (GraphPad Software). Unpaired two-tailed Student’s t-test was used to compare the means of two groups; one-way ANOVA was used with Tukey’s post-hoc test for pairwise comparison of means from more than two groups, or with Dunnett’s post-hoc test for comparison of means relative to the mean of a control group. Differences with a calculated *p*-value above 0.05 were considered non-significant and statistically significant differences were noted as follows: * = *p* < 0.05; ** = *p* < 0.01; *** = *p* < 0.001; **** = *p* <0,0001.

## Supplementary Materials

Supplementary Figure 1: Structure of *L. monocytogenes* RmlT and *S. aureus* TarS, TarP. Supplementary Figure 2: Amino-acid sequence alignment of RmlT, TarS and TarP. Supplementary Figure 3: Properties of RmlT. Supplementary Figure 4: Electron density map of the donor substrates TDP-rhamnose and UDP-glucose in different forms of RmlT. Supplementary Figure 5: Active site of RmlT with TDP-rhamnose or UDP-glucose. Supplementary Figure 6: Omit map of the donor substrate TDP-rhamnose in different RmlT copies. Supplementary Figure 7: Active sites of RmlT, TarS and TarP. Supplementary Figure 8: Active site of RmlT with and without TDP-rhamnose. Supplementary Figure 9: Omit map of the donor substrate TDP-rhamnose in different RmlT_D198A_ copies. Supplementary Figure 10: Omit map of the donor substrate UDP-glucose in different RmlT copies. Supplementary Figure 11: Omit map of 3-RboP in different copies of RmlT. Supplementary Figure 12: Omit map of the 4-RboP in different copies of RmlT. Supplementary Figure 13: HEPES binding sites. Supplementary Figure 14: Quantification of RmlT protein abundance in *L. monocytogenes* lysates. Supplementary Table 1: Quality of fit of protein chains to electron-density map. Supplementary Table 2: Bacterial strains and plasmids. Supplementary Table 3: Primer sequences.

## Author’s contributions

RM, JMC and DC conceptualize the overarching aims of the research study. RM, TBC, RP, TV, JDC-C, SS, JMC and DC conceived and designed the experiments. RM, TBC, TV and RP performed the experiments and data acquisition. RM, TBC, RP, SS, JMC and DC analysed and interpreted the data. JM-C and DC had management as well as coordination responsibility for the execution of the research work. JDC-C, JM-C and DC contributed to the acquisition of the financial supports and resources leading to this publication. All authors contributed to the drafting of some parts of the manuscript, including reading and revising critically the manuscript for important intellectual content, as well as approval of the final version.

## Acknowledgments

The work leading to these results was supported by FEDER – “Fundo Europeu de Desenvolvimento Regional” funds through the NORTE 2020 - Norte Portugal Regional Operational Programme, Portugal 2020, and by Portuguese funds through FCT - Fundação para a Ciência e a Tecnologia/Ministério da Ciência, Tecnologia e Ensino Superior in the framework of the project NORTE-01-0145-FEDER-030020 and PTDC/SAU-INF/30020/2017, and by the “la Caixa” Foundation and FCT, I.P. under project CI21-00035 and LCF/PR/ HR23-00682. TBC was supported by FCT (2021.01203.CEECIND).

The authors acknowledge the SOLEIL and ALBA synchrotrons for access and thank their staff for help with data collection. Support from the Biochemical and Biophysical Technologies, Animal Facility, Proteomics and X-ray Crystallography scientific platforms of i3S (Porto, Portugal) is also acknowledged. The authors extend their appreciation to Martin J. Loessner from ETH Zurich (Switzerland) that kindly provided the gp17-GFP recombinant construct.

