## supplementary material for "Molecular properties of RmlT, a wall teichoic acid rhamnosyltransferase that modulates virulence in *Listeria monocytogenes*"

**Supplementary Information**

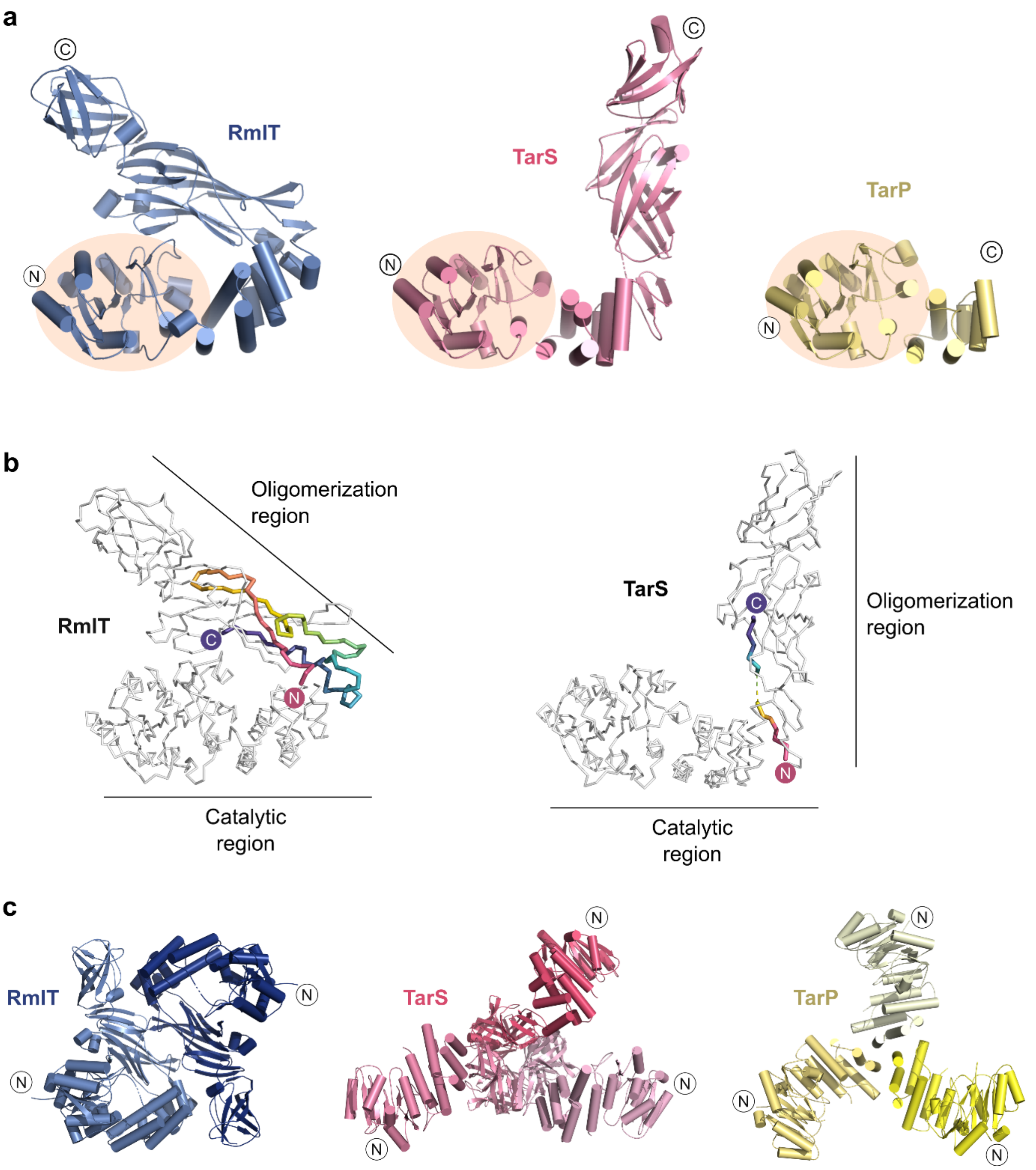


**Supplementary Figure 1:** Structures of *L. monocytogenes* RmlT and *S. aureus* TarS and TarP. **a)** Cartoon representation of subunits of RmlT (dark blue; PDB code 8BZ4), TarS (pink; PDB code 5TZ8) and TarP (yellow; PDB code 6H1J). N and C termini are indicated. Catalytic domains are highlighted by a wheat ellipse. **b)** Ribbon representation of RmlT (left) and TarS (right) with linker connecting the catalytic and the oligomerization regions colored from pink (catalytic region) to purple (oligomerization region). **c)** Cartoon representation of dimeric RmlT and trimeric TarS and TarP. Subunits are coloured in different shades. N termini are indicated.

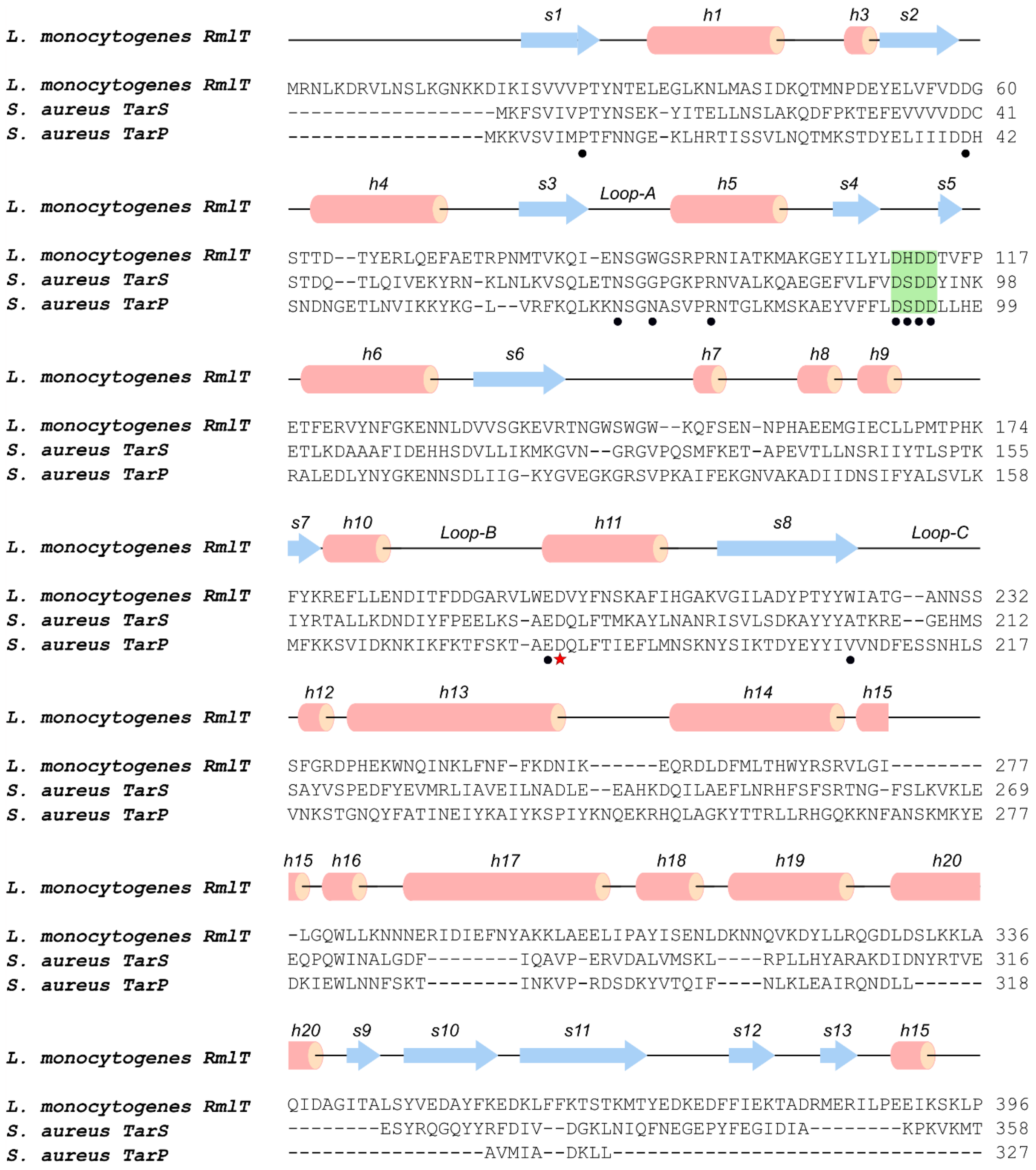


**Supplementary Figure 2:** Amino-acid sequence alignment of RmlT, TarS and TarP. Amino acid sequences of *L. monocytogenes* RmlT (Uniprot primary accession code Q8Y838), *S. aureus* TarS (Uniprot primary accession code A0A0H3JPC6) and *S. aureus* TarP (Uniprot primary accession code A0A0H3JNB0) were aligned with ClustalOmega. Alignment is included up to RmlT residue 396. Secondary structure elements of RmlT were identified with PROMOTIF and are indicated above the alignment. Putative catalytic residue is marked with a red star; other donor substrate-interacting residues are marked with black dot. DXDD motif involved in metal coordination is highlighted with a green square.

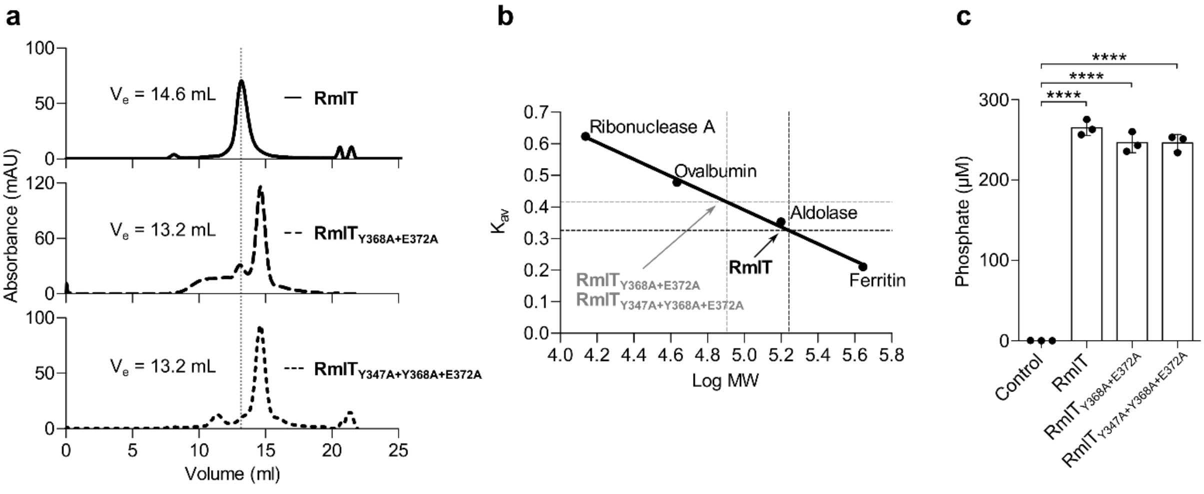


**Supplementary Figure 3:** Properties of RmlT. **a)** Analytical size-exclusion chromatogram of RmlT, RmlT$_{Y368A+E372A}$ and RmlT$_{Y347A+Y368A+E372A}$. Elution volumes were 13.2 mL for RmlT (dotted line) and 14.6 mL for RmlT$_{Y368A+E372A}$ and RmlT$_{Y347A+Y368A+E372A}$ in a Superdex 200 10/300 GL column. **b)** Calibration curve of column used in a). Ribonuclease A (13.7 kDa), Ovalbumin (43 kDa), Aldolase (158 kDa) and Ferritin (440 kDa) were used as standards. Wild-type RmlT (black dotted lines) eluted with an apparent molecular weight of 174 kDa, compatible with a dimer (theoretical MW = 143 kDa). Mutated variants (grey dotted line) eluted preferentially with an apparent molecular weight of 80 kDa, compatible with a monomer (theoretical MW = 71.5 kDa). **c)** Coupled functional assay quantifying the amount of phosphate released after hydrolysis of TDP molecules generated by RmlT. Reactions were performed for 60 minutes with 5 mM WTAs, 1 mM TDP-rhamnose and 10 µM protein. Control reaction was performed without protein. Mean ± SD (n = 3) individual measurements are shown; one-way ANOVA; **** $p < 0.0001$.

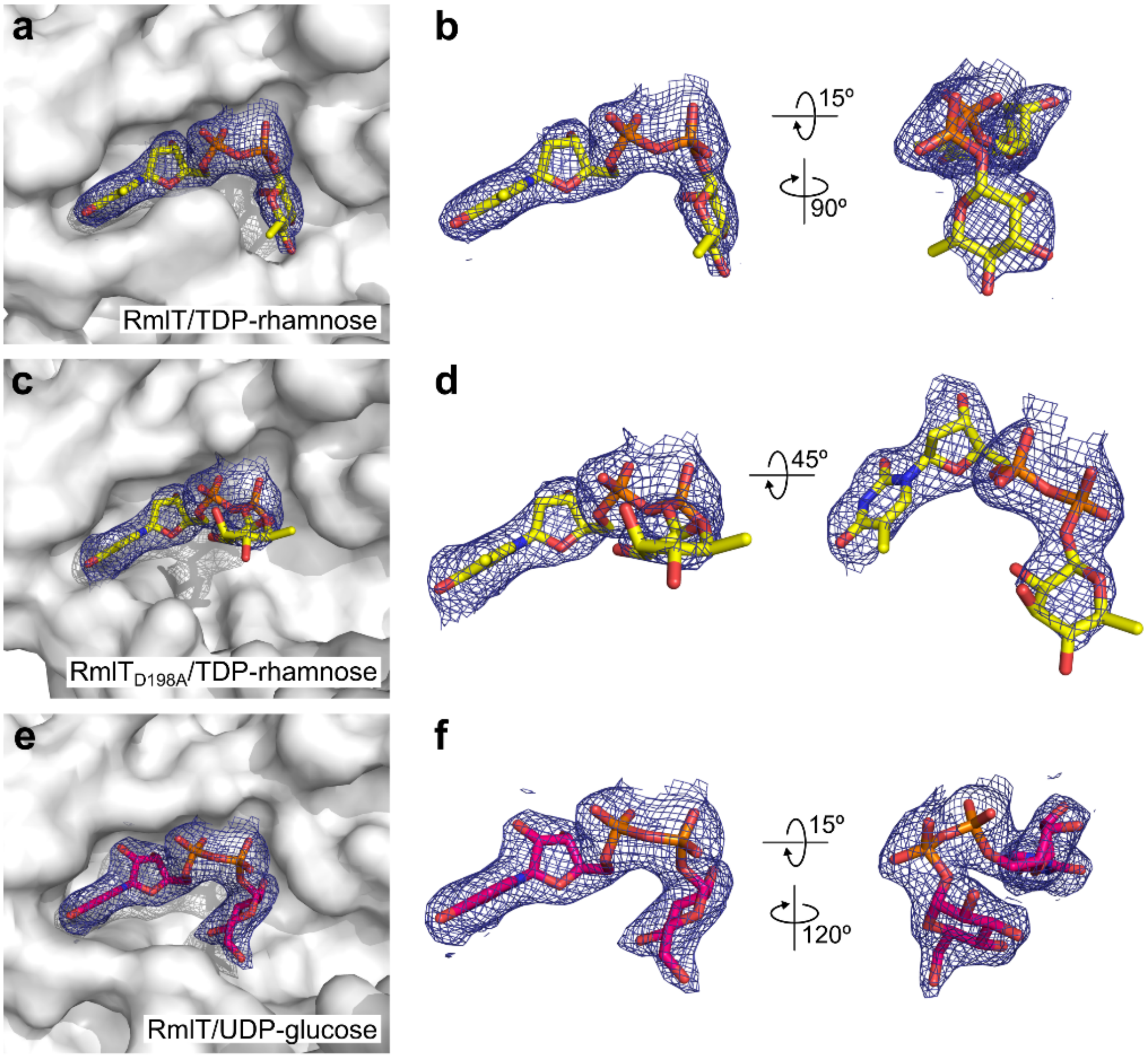


**Supplementary Figure 4:** Electron density map of the donor substrates TDP-rhamnose and UDP-glucose in different forms of RmlT. **a)**, **c)** and **e)** Surface representations of donor substrate binding pocket in wild-type (a and e) and D198A (c) RmlT proteins. TDP-rhamnose (a and c) and UDP-glucose (e) are represented as sticks and enveloped by the 2Fo-Fc electron density map contoured at 1.0 σ (blue mesh). **b)**, **d)** and **f)** Two different orientations of a zoomed view of TDP-rhamnose (b and d) and UDP-glucose (F) molecules and respective maps.

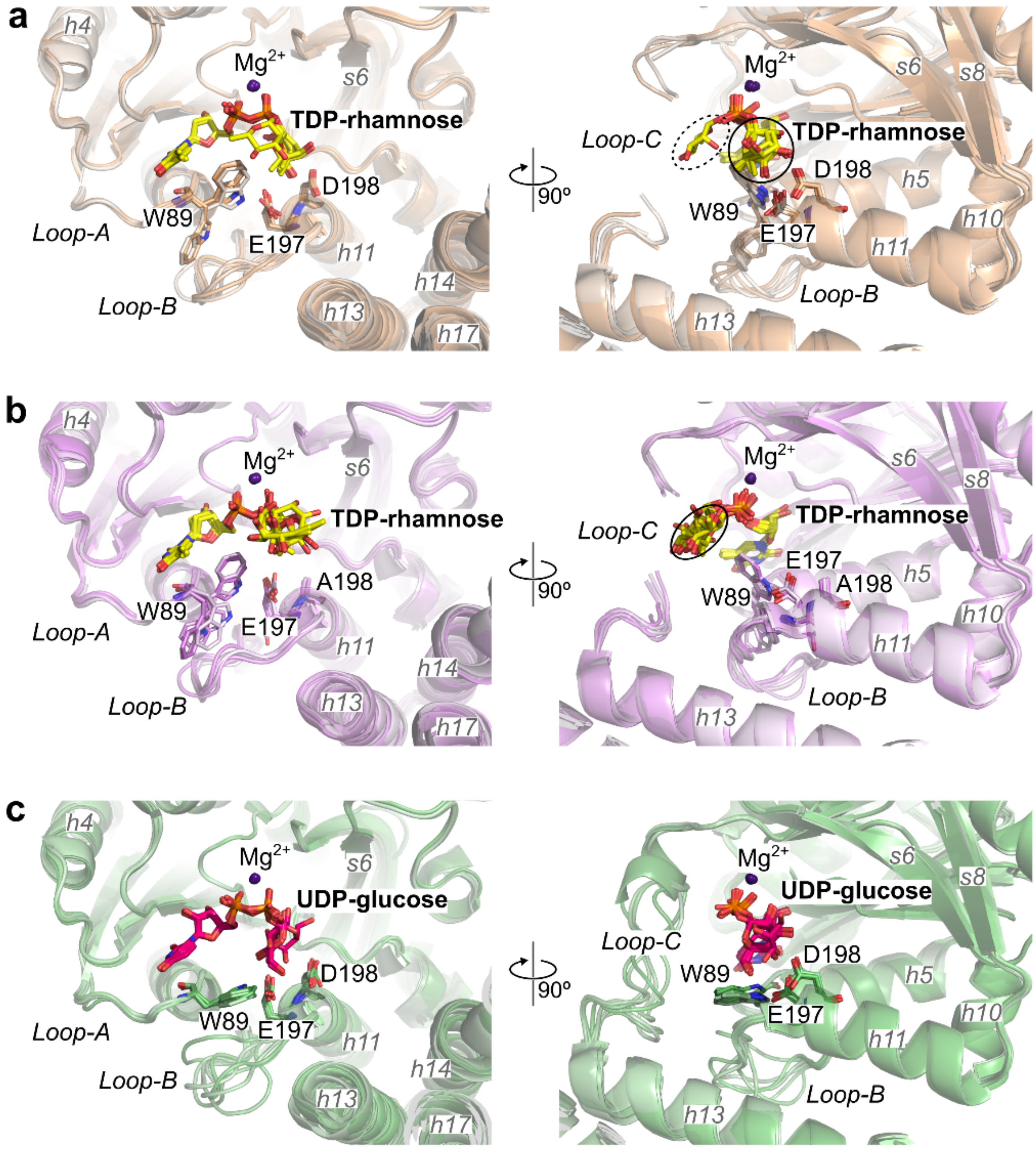


**Supplementary Figure 5:** Active site of RmlT with TDP-rhamnose or UDP-glucose. Superposition of the active-site of all RmlT (**a)** and **c)**) or RmlT$_{D198A}$ (**b)**) subunits present in the asymmetric unit and containing TDP-rhamnose (**a)** and **b)**) or UDP-glucose (**c)**). Left and right panels are related by a 90º rotation. Rhamnose moiety can adopt two positions in the active site of RmlT (a, right panel), highlighted by the circle and ellipse, with solid line indicating the position more frequently observed. In the RmlT$_{D198A}$ active site, all rhamnose moieties were found in the same position, which is indicated by the ellipse (b, right panel). Loop-C is completely modelled only in the RmlT/UDP-glucose complex (c). In a) and b), W89 displays more than one position. In c), W89 is well stabilized by contacts with E197 that, in turn, is bound to the glucose moiety of the donor substrate. Noteworthy, Loop-B, just before residues E197 and D198, adopts different conformations in RmlT/UDP-glucose but not in the

TDP-rhamnose structures. Superposition of the chains were performed through Cα pairs of the residues 110-114. RMSD relative to 8BZ7 chain D: 0.034 Å (8BZ7 chain A), 0.046 Å (8BZ7 chain B), 0.027 Å (8BZ7 chain C), 0.042 Å (8BZ7 chain E), 0.060 Å (8BZ7 chain F), 0.046 Å (8BZ8 chain A), 0.060 Å (8BZ8 chain B), 0.023 Å (8BZ8 chain C), 0.089 Å (8BZ8 chain D), 0.051 Å (8BZ8 chain E), 0.047 Å (8BZ8 chain F), 0.027 Å (8BZ6 chain A), 0.039 Å (8BZ6 chain B), 0.034 Å (8BZ6 chain C), 0.042 Å (8BZ6 chain D), 0.025 Å (8BZ6 chain E), 0.037 Å (8BZ6 chain F).

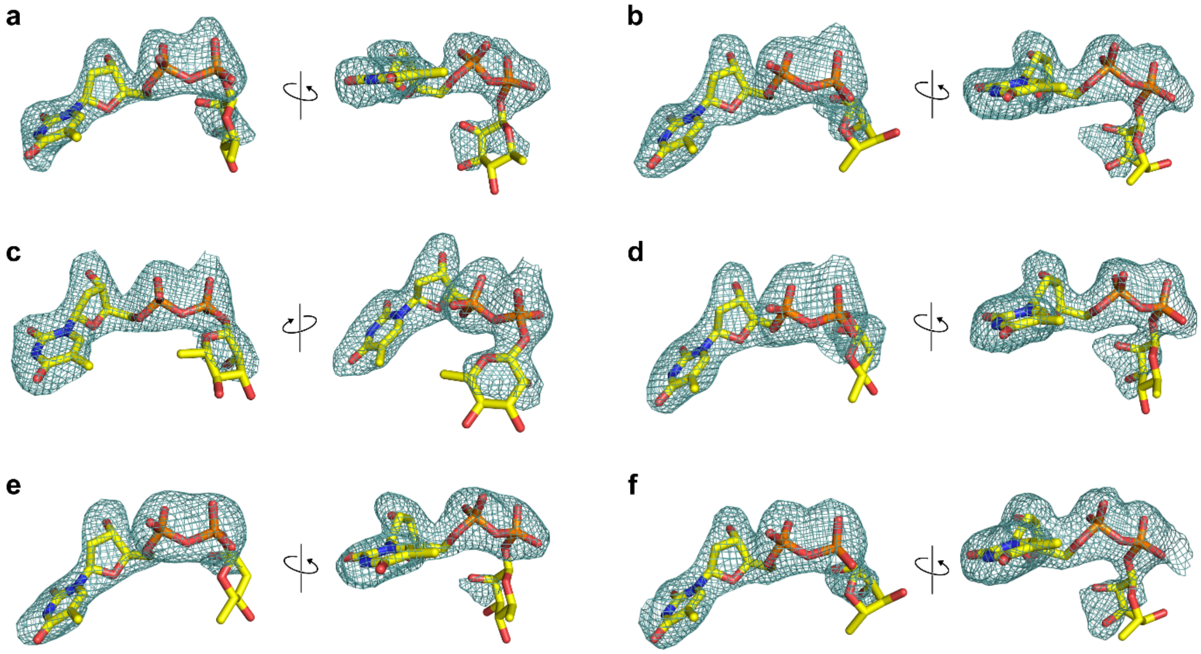


**Supplementary Figure 6:** Omit map of the donor substrate TDP-rhamnose in different RmlT copies. TDP-rhamnose molecules found in the active site of the RmlT subunits present in the asymmetric unit of PDB ID 8BZ7, represented as sticks and shown in two different positions, are enveloped by corresponding omit map contoured at 3.0 σ (teal mesh). Although well-defined in the TDP moiety, omit map shows poor quality in the sugar. Left and right panels are related by rotation. **a)** chain A, **b)** chain B, **c)** chain C, **d)** chain D, **e)** chain E, **f)** chain F.

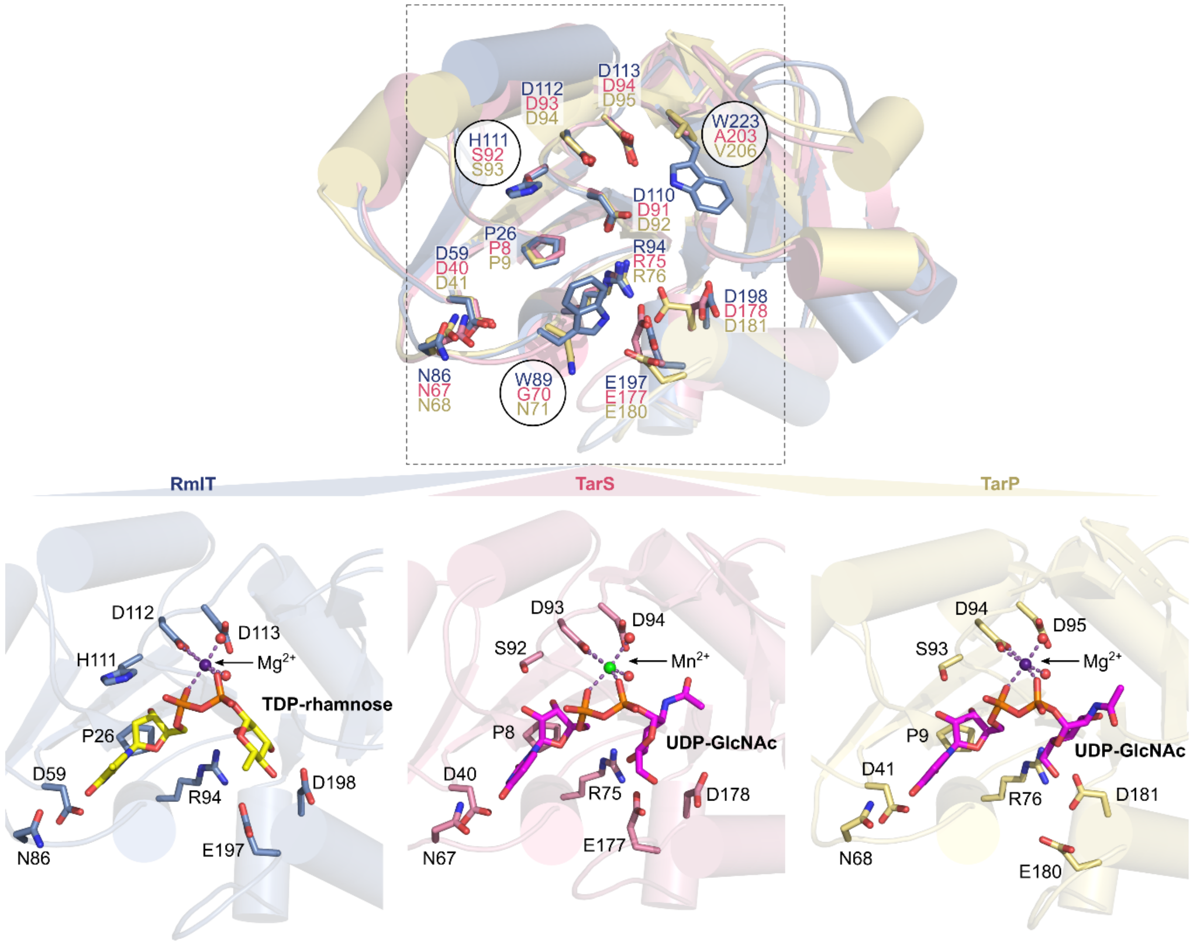


**Supplementary Figure 7:** Active sites of RmlT, TarS and TarP. Top panel, Cartoon representation of superposed active sites of *L. monocytogenes* RmlT (blue; PDB code 8BZ7, chain D) and *S. aureus* TarS (pink; PDB code 5TZE, chain C) and TarP (yellow; PDB code 6H2N, chain B). Superposition was performed through Cα of the residues shown as sticks in top panel (RMSD of 0.831 for TarS and 0.663 for TarP). Donor substrate interacting residues are represented as sticks. Amino acid differences between proteins are highlighted by black circles. Bottom panel, Cartoon representation of the active site region of RmlT (blue), TarS (pink) and TarP (yellow) with conserved residues (excepting RmlT D110, TarS D91 and TarP D92) represented as sticks. TDP-rhamnose (yellow, RmlT) and UDP-GlcNAc (cyan, TarS and TarP) are shown as sticks. $Mg^{2+}$ (purple), $Mn^{2+}$ (green) and water (red) molecules are shown as spheres and metal coordination is represented by dashed light purple lines.

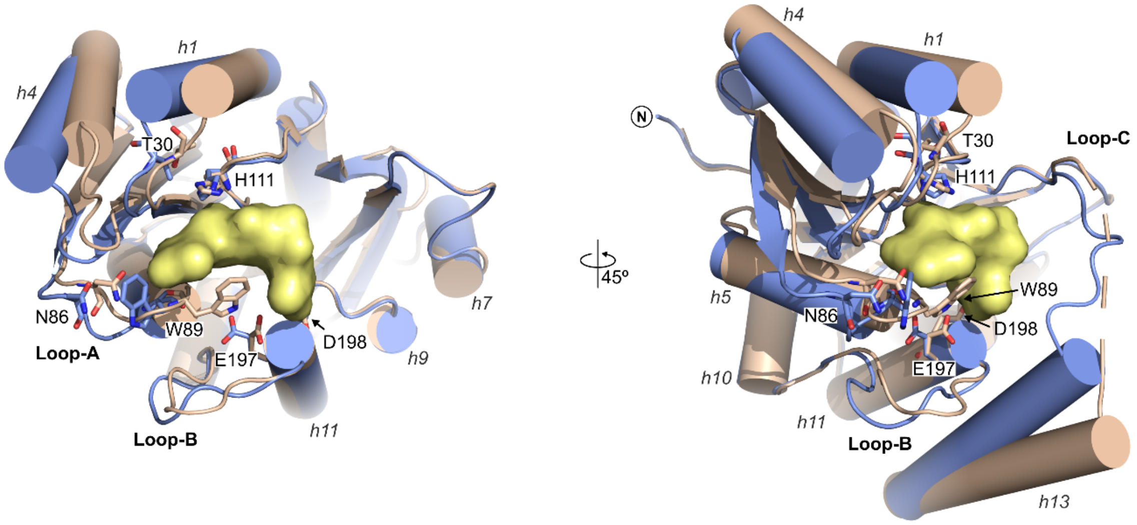


**Supplementary Figure 8:** Active site of RmlT with and without TDP-rhamnose. Cartoon representation of the active site of RmlT with (wheat; PDB code 8BZ7, chain D) and without (blue; PDB code 8BZ5, chain C) TDP-rhamnose (yellow surface representation). Residues involved in substrate binding are represented as sticks. Loops, helices and N terminus are indicated. Loop-C and helix h13 were hidden in left view. Structures were superposed using 5 Cα atoms pairs (residues 110-114) with RMSD of 0.198. Panels are related by rotation.

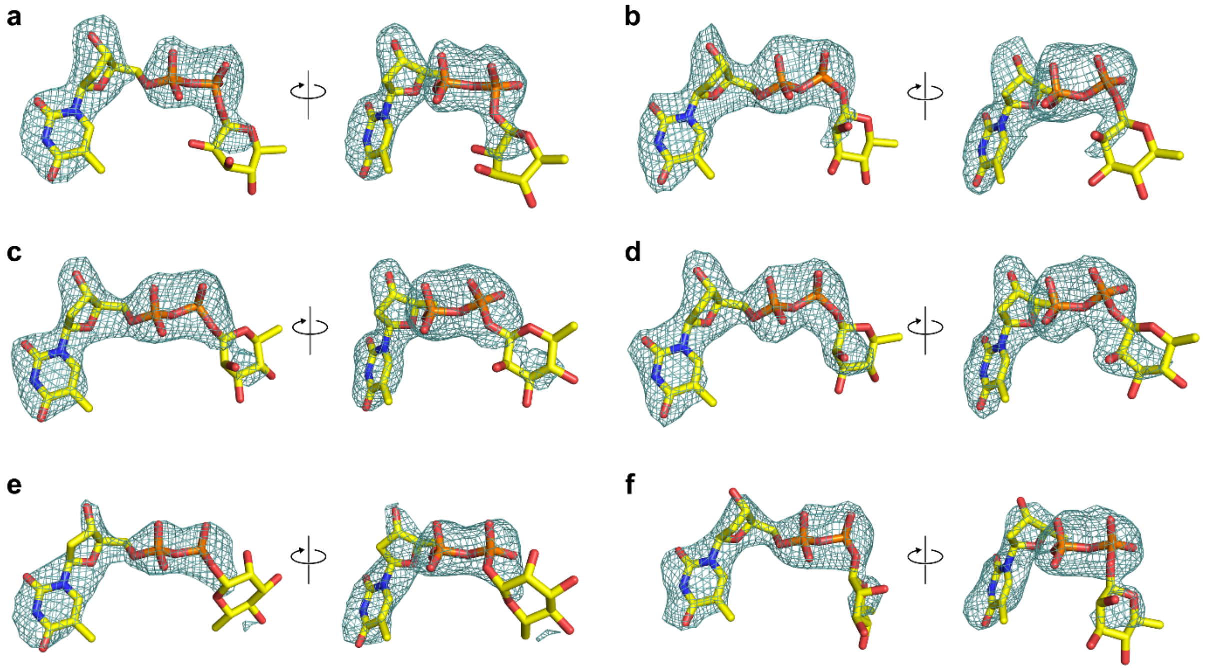


**Supplementary Figure 9:** Omit map of the donor substrate TDP-rhamnose in different $RmlT_{D198A}$ copies. TDP-rhamnose molecules found in the active site of the $RmlT_{D198A}$ subunits present in the asymmetric unit (PDB code 8BZ8), represented as sticks and shown in two different positions, are enveloped by corresponding omit map contoured at 3.0 σ (teal mesh). Omit map show low definition at the sugar region of the donor substrate but clearly restrict its relative position. Left and right panels are related by rotation. **a)** chain A, **b)** chain B, c) chain C, **d**) chain D, **e)** chain E, **f)** chain F.

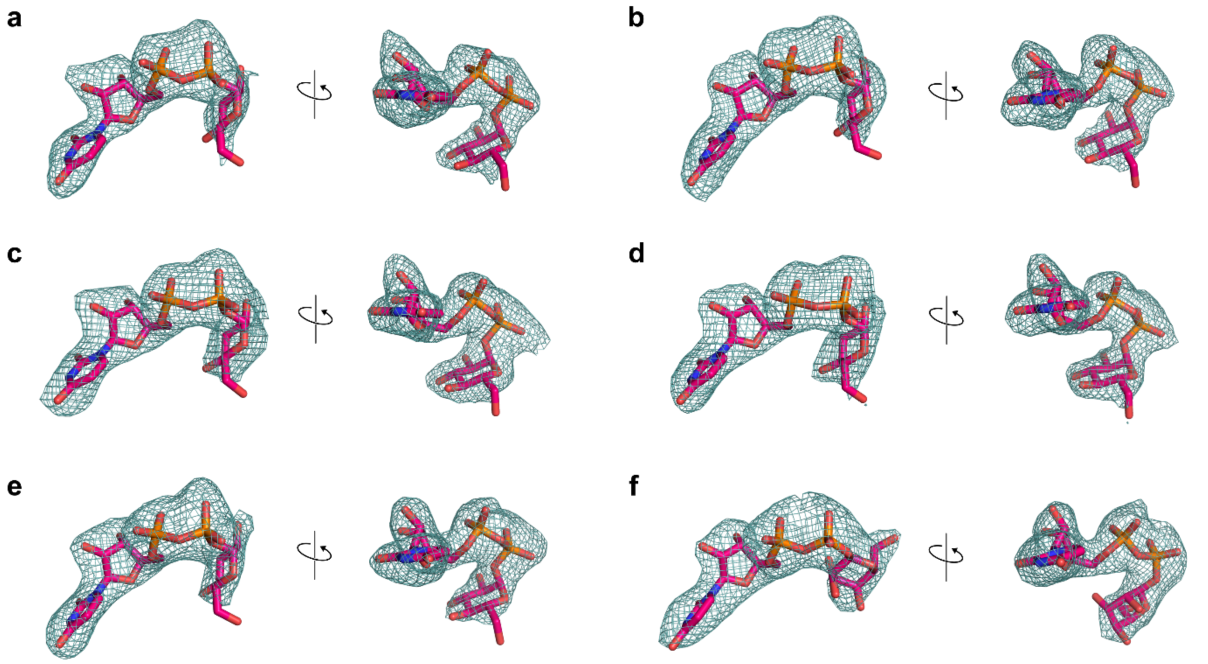


**Supplementary Figure 10:** Omit map of the donor substrate UDP-glucose in different RmlT copies. UDP-glucose molecules found in the active site of the RmlT subunits present in the asymmetric unit (PDB Icode 8BZ6), represented as sticks and shown in two different positions, are enveloped by corresponding omit map contoured at 3.0 σ (teal mesh). Map is well defined for the entire substrate molecule. Left and right panels are related by rotation. **a)** chain A, **b)** chain B, **c)** chain C, **d)** chain D, **e)** chain E, **f)** chain F.

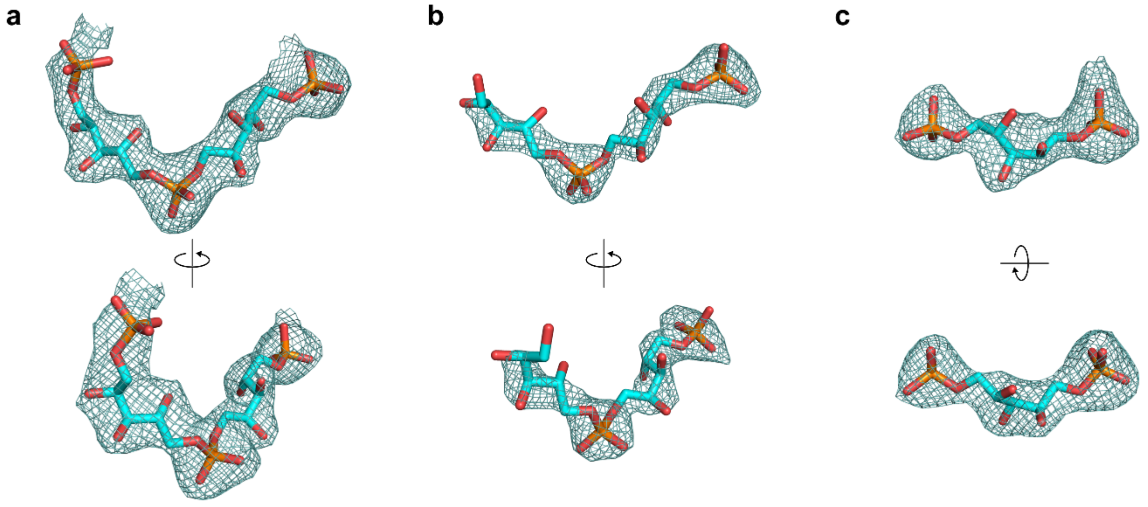


**Supplementary Figure 11:** Omit map of 3-RboP in different copies of RmlT. RboP molecules found in RmlT subunits present in the asymmetric unit (PDB code 9GZJ), represented as sticks and shown in two different positions, are enveloped by corresponding omit map contoured at 3.0 σ (teal mesh). Top and bottom panels are related by rotation. RboP molecules in binding site 1 of chain A (**a**) and chain B (**b**) and in the binding site 2 of chain A (**c**).

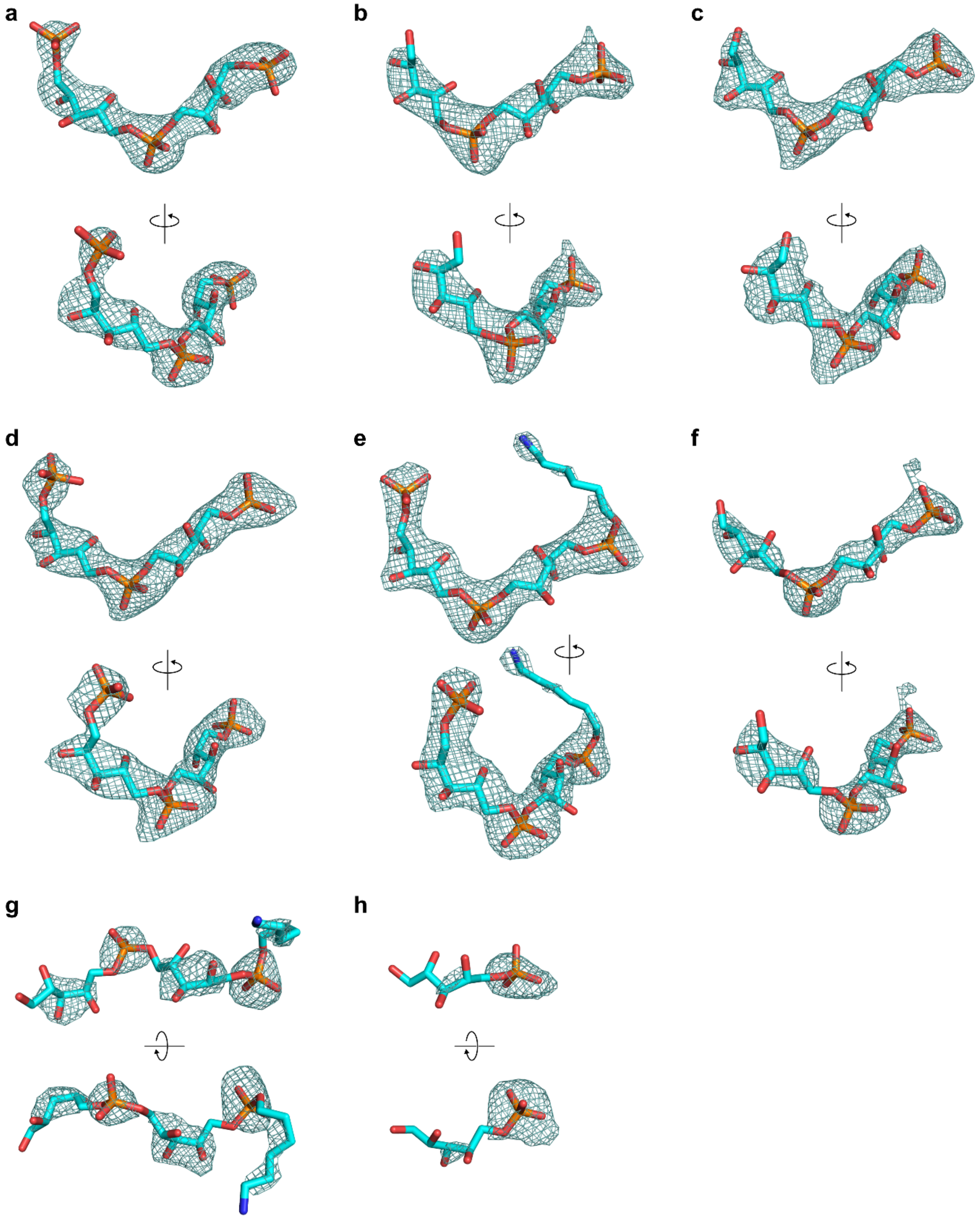


**Supplementary Figure 12:** Omit map of the 4-RboP in different copies of RmlT. RboP molecules found in RmlT subunits present in the asymmetric unit (PDB code 9GZK), represented as sticks and shown in two different positions, are enveloped by corresponding omit map contoured at 3.0 σ (teal mesh). Top and bottom panels are related by rotation. e RboP in binding site 1 shown in **a)** chain A, **b)** chain B, **c)** chain C, **d)** chain D, **e)** chain E, **f)** chain F. RboP in binding site 2 shown in **g)** chain B, **h)** chain F.

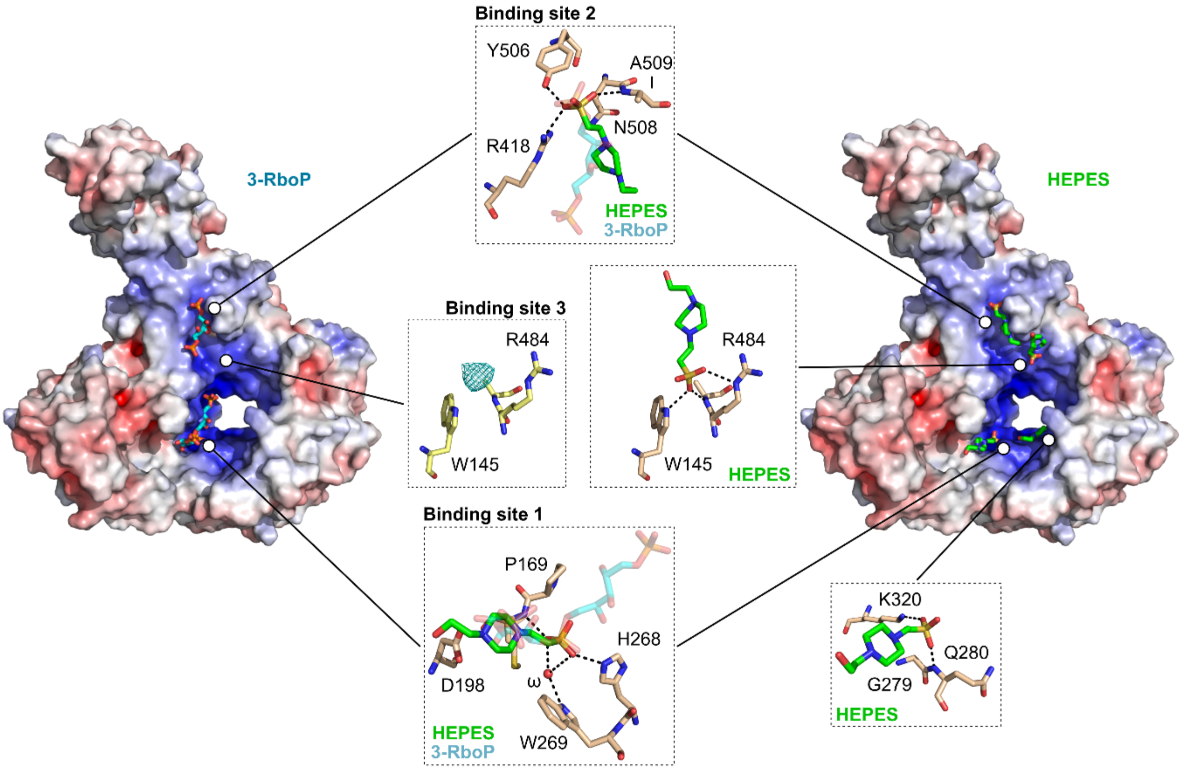


**Supplementary Figure 13:** HEPES binding sites. Electrostatic surface representation of RmlT subunit bound to: Left, RboP molecules (PDB code 9GZJ, chain A); Right, HEPES (PDB code 8BZ5, chain B). Proteins are shown in same orientation after superposition Cα of resides 238-516 (RMSD = 0.378 Å). Details of the binding sites containing HEPES (in green stick) are shown. RboP chains found in the same sites as HEPES are shown in cyan with transparency. Strong electron-density (teal mesh countered at 3.0 σ) compatible with a phosphate group is present in binding site 3 of RmlT/3-RboP. Hydrogen bonds are represented as black dashes lines.

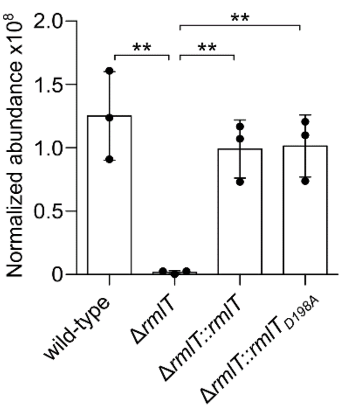


**Supplementary Figure 14:** Quantification of RmlT protein abundance in *L. monocytogenes* lysates. RmlT abundance in *L. monocytogenes* EGDe (wild-type), *L. monocytogenes* EGDe Δ*rmlT*, *L. monocytogenes* EGDe Δ*rmlT::rmlT* or *L. monocytogenes* EGDe Δ*rmlT::rmlT*$_{D198A}$ was quantified by mass spectrometry. Abundancy values were normalized between samples by total peptide content. Mean ± SD (n = 3) and individual measurements are shown; one-way ANOVA; ** $p < 0.001$.

**Supplementary Table 1:** Quality of fit of protein chains to electron-density map.

| PDB code | Chain | Number of residues built (% out of 624) | <RSRZ> | RSRZ outliers #/% | Median occupancy-weighted average B-factor (Å$^2$) |
|---|---|---|---|---|---|
| 8BZ4 | A | 595 (95%) | 0.04 | 3 / 0% | 49 |
| | B | 566 (90%) | 1.00 | 104 / 18% | 90 |
| 8BZ5 | A | 603 (96%) | 0.13 | 33 / 5% | 52 |
| | B | 600 (96%) | 0.03 | 15 / 2% | 52 |
| | C | 608 (97%) | 0.29 | 34 / 5% | 58 |
| | D | 598 (95%) | 0.23 | 36 / 6% | 54 |
| 8BZ6 | A | 606 (97%) | 0.24 | 10 / 1% | 61 |
| | B | 605 (96%) | 0.71 | 78 / 12% | 72 |
| | C | 606 (97%) | 0.31 | 13 / 2% | 63 |
| | D | 606 (97%) | 0.23 | 11 / 1% | 61 |
| | E | 606 (97%) | 0.15 | 7 / 1% | 61 |
| | F | 606 (97%) | 0.62 | 54 / 8% | 75 |
| 8BZ7 | A | 601 (96%) | 0.12 | 9 / 1% | 48 |
| | B | 601 (96%) | 0.18 | 8 / 1% | 51 |
| | C | 600 (96%) | 0.10 | 12 / 2% | 51 |
| | D | 599 (95%) | 0.48 | 58 / 9% | 54 |
| | E | 598 (95%) | 0.16 | 12 / 2% | 59 |
| | F | 598 (95%) | 0.60 | 66 / 11% | 64 |
| 8BZ8 | A | 603 (96%) | -0.08 | 4 / 0% | 55 |
| | B | 601 (96%) | 0.31 | 36 / 5% | 66 |
| | C | 603 (96%) | 0.04 | 4 / 0% | 53 |
| | D | 603 (96%) | -0.01 | 4 / 0% | 54 |
| | E | 599 (95%) | 0.05 | 7 / 1% | 64 |
| | F | 599 (95%) | 0.39 | 50 / 8% | 71 |
| 9GZJ | A | 608 (97%) | -0.32 | 2 / 0% | 54 |
| | B | 573 (91%) | 0.54 | 28 / 4% | 92 |
| 9GZK | A | 600 (96%) | -0.28 | 5 / 0% | 63 |
| | B | 601 (96%) | 0.08 | 11 / 1% | 81 |
| | C | 599 (95%) | -0.17 | 2 / 0% | 67 |
| | D | 599 (95%) | -0.07 | 1 / 0% | 75 |
| | E | 597 (95%) | -0.25 | 4 / 0% | 64 |
| | F | 544 (87%) | 0.53 | 30 / 5% | 98 |

RSRZ is a normalized Real Space R-factor that reflects the quality of fit between a residue and the electron-density map.

**Supplementary Table 2**: Bacterial strains and plasmids.

| Plasmid/Strain | Description | Source |
|---|---|---|
| **Strains** | | |
| *L. monocytogenes* EGDe | *Listeria monocytogenes* EGD-e (serotype 1/2a). | [51] |
| *L. monocytogenes* EGDe Δ*rmlT* | Isogenic mutant of *Listeria monocytogenes* EGD-e lacking the gene *lmo1080 (= rmlT)*. | [13] |
| *L. monocytogenes* EGDe Δ*rmlT::rmlT* | Isogenic mutant of *Listeria monocytogenes* EGD-e lacking the gene *lmo1080 (= rmlT),* complemented with the gene *lmo1080* (= *rmlT*). | [13] |
| *L. monocytogenes* EGDe Δ*lmo1079* | Isogenic mutant of *Listeria monocytogenes* EGD-e lacking the gene *lmo1079*. | This study |
| *L. monocytogenes* EGDe Δ*rmlT*Δ*lmo1079* | Isogenic mutant of *Listeria monocytogenes* EGD-e lacking the genes *lmo1080 (= rmlT)* and *lmo1079*. | This study |
| *L. monocytogenes* EGDe Δ*rmlT::rmlT*$_{Y368A+E372A}$ | Isogenic mutant of *Listeria monocytogenes* EGD-e lacking the gene *lmo1080 (= rmlT),* complemented with the gene *lmo1080* (= *rmlT*) mutated for Y368 and E372. | This study |
| *L. monocytogenes* EGDe Δ*rmlT::rmlT*$_{Y347A+Y368A+E372A}$ | Isogenic mutant of *Listeria monocytogenes* EGD-e lacking the gene *lmo1080 (= rmlT),* complemented with the gene *lmo1080* (= *rmlT*) mutated for Y347, Y368 and E372. | This study |
| *L. monocytogenes* EGDe Δ*rmlT::rmlT*$_{D198A}$ | Isogenic mutant of *Listeria monocytogenes* EGD-e lacking the gene *lmo1080 (= rmlT),* complemented with the gene *lmo1080* (= *rmlT*) mutated for D198. | This study |
| *L. monocytogenes* EGDe Δ*rmlT::rmlT*$_{E197A}$ | Isogenic mutant of *Listeria monocytogenes* EGD-e lacking the gene *lmo1080 (= rmlT),* complemented with the gene *lmo1080* (= *rmlT*) mutated for E197. | This study |
| *L. monocytogenes* EGDe Δ*rmlT::rmlT*$_{W223A}$ | Isogenic mutant of *Listeria monocytogenes* EGD-e lacking the gene *lmo1080 (rmlT),* complemented with the gene *lmo1080* (*rmlT*) mutated for W223. | This study |
| *L. monocytogenes* EGDe Δ*rmlT::rmlT*$_{W89A}$ | Isogenic mutant of *Listeria monocytogenes* EGD-e lacking the gene *lmo1080* (= *rmlT),* complemented with the gene *lmo1080* (= *rmlT*) mutated for W89. | This study |
| *E. coli* DH5α | Strain engineered to maximize transformation efficiency; recA1$_{G160D}$ endA1$^-$ *lacZ*ΔM15. | Novagen |
| *E. coli* B834(DE3) | Methionine auxotroph for labeling proteins with selenomethionine for crystallography. | Novagen |
| *E. coli* BL-21 (DE3) | Cells for protein expression; F$^-$ *ompT hsdS*$_B$ ($r_B^- m_B^-$) gal dcm (DE3). | Novagen |
| **Plasmids** | | |
| pET28b | Vector for cloning and expression of recombinant proteins in *E. coli*. | Novagen |

| | | |
|---|---|---|
| pET28b-*rmlT* | Vector for inducible expression of recombinant RmlT. | This study |
| pET28b-*rmlT*$_{Y368A+E372A}$ | Vector for inducible expression of recombinant mutated RmlT. | This study |
| pET28b-*rmlT*$_{Y347A+Y368A+E372A}$ | Vector for inducible expression of recombinant mutated RmlT. | This study |
| pET28b-*rmlT*$_{D198A}$ | Vector for inducible expression of recombinant mutated RmlT. | This study |
| pET28b-*rmlT*$_{E197A}$ | Vector for inducible expression of recombinant mutated RmlT. | This study |
| pET28b-*rmlT*$_{W223A}$ | Vector for inducible expression of recombinant mutated RmlT. | This study |
| pET28b-*rmlT*$_{W89A}$ | Vector for inducible expression of recombinant mutated RmlT. | This study |
| pET28b-*rmlT*$_{W145C+A472C}$ | Vector for inducible expression of recombinant mutated RmlT. | This study |
| pPL2 | *L. monocytogenes* conjugative vector. | [52] |
| pPL2-*rmlT*$_{Y368A+E372A}$ | Integrative plasmid expressing *rmlT*$_{Y368A+E372A}$. | This study |
| pPL2-*rmlT*$_{Y347A+Y368A+E372A}$ | Integrative plasmid expressing *rmlT*$_{Y347A+Y368A+E372A}$. | This study |
| pPL2-*rmlT*$_{D198A}$ | Integrative plasmid expressing *rmlT*$_{D198A}$. | This study |
| pPL2-*rmlT*$_{E197A}$ | Integrative plasmid expressing *rmlT*$_{E197A}$. | This study |
| pPL2-*rmlT*$_{W223A}$ | Integrative plasmid expressing *rmlT*$_{W223A}$. | This study |
| pPL2-*rmlT*$_{W89A}$ | Integrative plasmid expressing *rmlT*$_{W89A}$. | This study |

**Supplementary Table 3**: Primer sequences.

| | Name | Sequence (5' – 3') |
|---|---|---|
| **Cytoplasmic overexpression of recombinant proteins** | | |
| #1 | RmlT Fwd_N-terminal HisTag | GAAAACCTGTACTTCCAGGGCATGAGAAATTTAAAAGATAGAG |
| #2 | RmlT Rev_N-terminal HisTag | GTTAGCAGCCGGATCTCAAAGCGTTATTTTAATTTGCTG |
| #3 | pET28b Fwd_N-terminal HisTag | TGAGATCCGGCTGCTAACAAAGCCCGAAAGGAAGCTGAG |
| #4 | pET28b Rev_N-terminal HisTag | GCCCTGGAAGTACAGGTTTTCGTGATGATGATGATGATGGCTGCTG |
| **Site-directed mutagenesis** | | |
| #5 | RmlT_D198A_Fwd | CGAGTTCTTTGGGAAGCAGTATATTTTAAC |
| #6 | RmlT_D198A_Rev | GTTAAAATATACTGCTTCCCAAAGAACTCG |
| #7 | RmlT_E197A_Fwd | GCTCGAGTTCTTTGGGCAGATGTATATTTTAAC |
| #8 | RmlT_E197A_Rev | GTTAAAATATACATCTGCCCAAAGAACTCGAGC |
| #9 | RmlT_W89A_Fwd | GAAAACTCTGGTGCAGGAAGCAGACC |
| #10 | RmlT_W89A_Rev | GGTCTGCTTCCTGCACCAGAGTTTTC |
| #11 | RmlT_W223A_Fwd | CCTACGTACTATGCAATTGCAACTGGTGC |
| #12 | RmlT_W223A_Rev | GCACCAGTTGCAATTGCATAGTACGTAGG |
| #13 | RmlT_E372A+Y368A_Fwd | GTACAAAAATGACAGCAGAAGACAAAGCAGATTTCTTTATTG |
| #14 | RmlT_E372A+Y368A_Rev | CAATAAAGAAATCTGCTTTGTCTTCTGCTGTCATTTTTGTAC |
| #15 | RmlT_Y347A_Fwd | GGAATTACAGCGCTTAGCGCAGTGGAAGATGCTTAC |
| #16 | RmlT_Y347A_Rev | GTAAGCATCTTCCACTGCGCTAAGCGCTGTAATTCC |
| #17 | RmlT_W145C_Fwd | GAAGTAAGAACAAATGGTTGTTCATGGGGATGGAAG |
| #18 | RmlT_W145C_Rev | CTTCCATCCCCATGAACAACCATTTGTTCTTACTTC |
| #19 | RmlT_A472C_Fwd | CAACCATGGGATATTTGTACACGCTTTACCGGC |
| #20 | RmlT_A472C_Rev | GCCGGTAAAGCGTGTACAAATATCCCATGGTTG |
